# The rapid delivery of RNase H variants reveals distinct classes of RNA:DNA hybrids

**DOI:** 10.64898/2026.09.23.753687

**Authors:** Yara Bawadi, Amandine Basille, Yea-Lih Lin, Connor Kirke, Clara Bourgeay, David Cluet, Laetitia Vachez, Virgile Dervieux, Laura Guiguettaz, Pascal Bernard, Stephan Hamperl, Philippe Pasero, Emiliano P. Ricci, Vincent Vanoosthuyse

**Affiliations:** Université de Lyon, ENS de Lyon, Université Claude Bernard, CNRS UMR5239, Laboratoire de Biologie et Modélisation de la Cellule, Lyon, France; IGH, Univ Montpellier, CNRS, Montpellier, France; Institute of Epigenetics and Stem Cells, Helmholtz Center Munich, Munich, Germany; MRC Centre for Medical Mycology, University of Exeter, Exeter, UK

**Author notes:** equal contribution (Yara Bawadi, Amandine Basille).

**Keywords:** R-loops, RNase H1, RNA:DNA, Replication forks, Virus-Like Particles

## Abstract

Multiple lines of evidence suggest that RNase H1-sensitive RNA:DNA hybrids (thereafter referred to as ‘hybrids’) could disrupt the stability and restart of stalled replication forks. The prolonged over-expression of RNase H1 is the most popular method to manipulate hybrids *in situ* but it cannot discriminate between direct and indirect effects resulting from their loss. To overcome this limitation, we developed *iCoRD* (in Cellulo RNase H Delivery) to rapidly deliver comparable amounts of either *Escherichia coli* RNase H1 (RnhA) or control proteins in live cells, with a negligible impact on the transcriptome. *iCoRD*-delivered RnhA recognises hybrids *in situ*, is enriched in the vicinity of active replication forks and rescues replication fork progression in different stress conditions. While *iCoRD*-delivered RnhA had a clear impact on stress-induced, fork-proximal hybrids and R-ChIP signals, it had no effect on DRIP or S9.6 Cut&Tag signals, even when these were induced by stress. Our results strongly suggest the existence of distinct populations of hybrids that are best mapped by different methods and are differently sensitive to RNase H1 activity *in situ*.

## INTRODUCTION

In differentiated cells, early DNA replication forks progress slower and pause more frequently than late replication forks. Strikingly, drug treatments inhibiting gene transcription speed up early forks, suggesting that active transcription in early S phase hinders fork progression (van den Berg *et al*, 2024; Kurashima *et al*, 2024). A large body of evidence now supports the idea that conflicts between transcription and replication are a threat to chromosome integrity and that cells have evolved a number of mechanisms to either prevent or mitigate the consequences of such Transcription-Replication Conflicts (TRCs) (reviewed in (Browning & Merrikh, 2024; Uruci *et al*, 2026)). As TRCs might be more frequent in some tumours, to target the molecular response to TRCs could constitute a promising therapeutic strategy (Petropoulos *et al*, 2024; Gu *et al*, 2023; Patel *et al*, 2023; Dutrieux *et al*, 2021).

The reasons why active transcription threatens advancing replication forks are likely multifactorial. Both transcription and replication alter DNA topology, and excessive positive topological stress could mechanically impede the movement of replication forks by impairing DNA unwinding, particularly in the case of head-on conflicts between transcription and replication (reviewed in (Keszthelyi *et al*, 2016)). Altered DNA topology also increases the likelihood of co-transcriptional R-loops, three-stranded chromatin structures that form in chromatin when the nascent RNA winds around its DNA template resulting in a RNA:DNA hybrid and an unpaired non-template DNA strand (Stolz *et al*, 2019; Drolet, 2006). In support of the idea that excessive R-loop formation might impede the movement of replication forks and compromise genome integrity, a large body of evidence consistently shows that the adverse consequences of TRCs can be largely alleviated by over-expressing RNase H1, an enzyme that degrades the RNA moiety of RNA:DNA hybrids (reviewed in (Brickner *et al*, 2022)). Conversely, loss of RNase H1 results in the complete collapse of DNA replication at a single head-on TRC locus engineered in the genome of *Bacillus subtilis* (Lang *et al*, 2017; Lang & Merrikh, 2021). These concordant observations strongly suggest that RNA:DNA hybrids could interfere with faithful DNA replication and the resolution of TRCs. For convenience, ‘RNA:DNA hybrids’ will often be abbreviated to ‘hybrids’ in the following.

The population of hybrids that is sensitive to the over-expression of RNase H1 and whose loss mitigates the effect of replication stress on fork progression remains debated. Three different types of hybrids have so far been proposed to impinge on replication forks upon stress: (i) pre-existing, co-transcriptional R-loops might interfere with the progression of replication forks; (ii) post-replicative hybrids embedded in nascent DNA strands might form behind replication forks in response to TRCs and interfere with post-replicative processes important for fork stability and restart (Stoy *et al*, 2023; Heuzé *et al*, 2023; Xu *et al*, 2025; Song *et al*, 2025; Meroni *et al*, 2019). Whether these post-replicative hybrids are by-products of pre-existing R-loops that have been transferred behind forks by stress-induced DNA transactions or whether they are generated *de novo* in response to replication stress remains unclear; (iii) observations made in fission yeast suggested that transcription-independent but primase-dependent RNA molecules embedded in reversed forks could prevent the untimely resection of nascent DNA strands and thereby promote timely replication resumption (Audoynaud *et al*, 2023). Taken together, these observations suggest the existence of a complex, multi-origin and under-appreciated RNA:DNA landscape around stressed replication forks that could regulate the resection of nascent strands upon stress. Whether these different types of hybrids can all be efficiently eliminated by the over-expression of RNase H1 has not yet been addressed.

Another question is to understand the direct impact of hybrids on fork transactions upon replication stress. Currently, the most popular tool to manipulate hybrid levels in live cells is the prolonged overexpression of RNase H1 in asynchronous cell populations. However, this approach cannot discriminate between the direct and indirect effects of hybrid loss. Moreover, as R-loops have been proposed to be important transcription regulators (reviewed in (Niehrs & Luke, 2020)), the long-term over-expression of RNase H1 could significantly alter the transcriptome in a way that only indirectly mitigates the consequences of TRCs (reviewed in (Chédin *et al*, 2021)). The lengthy over-expression of RNase H1 therefore appears ill-adapted to the functional study of fork-proximal, stress-induced hybrids.

To circumvent this issue, we built an alternative strategy to manipulate and detect hybrids in live human cells. We used Virus-Like Particles (VLPs) to rapidly deliver a large amount of ready-made, catalytically active or inactive *Escherichia coli* RNase H1 (RnhA) variants. Importantly, this did not substantially impact the steady-state transcriptome. We provide compelling evidence that VLP-delivered RnhA recognises hybrids in live cells. It is also enriched in the vicinity of active replication forks and mitigates the impact of replication stress on replication fork progression. Consistent with this, we provide evidence that VLP-delivered RnhA is active against stress-induced, fork-proximal hybrids in *-cis*. However, it had no impact on DRIP or S9.6 Cut&Tag signals, two methods commonly used to map R-loops, even in stressed conditions where DRIP signals were strongly increased. On the contrary, VLP-delivered RnhA could eliminate R-ChIP signals at canonical loci. Our results show that, contrary to common expectations, not all types of RNA:DNA hybrids are eliminated by increasing the levels of RNase H1 in cells.

## RESULTS

### Rapid and flexible delivery of ready-made RNase H1 in the nuclei of live cells

We wondered whether Virus-Like Particles (VLPs), which have previously been used to deliver ready-made genome editors such as Cas9 in live vertebrate cells (see for example (Banskota *et al*, 2022; Mangeot *et al*, 2019)), could also be used to rapidly deliver ready-made sensors and regulators of RNA:DNA hybrids. VLPs are produced by the multimerization of the retroviral Gag structural polyprotein at the plasma membrane, which, when expressed alone or as a Gag-Pol fusion, induces the budding of vesicles lacking genetic information, and their subsequent release into the medium. Inclusion of viral envelope glycoproteins at their surface allows VLPs to fuse with target cells and deliver their protein cargo (Fig 1A). The fusion of a protein of interest to the C-terminus of Gag results in its inclusion within VLPs. A cleavage site for the viral protease, which is encoded within the Pol moiety of Gag-Pol, is inserted at the junction between Gag and the protein of interest to ensure its delivery into target cells as a separate entity (Fig 1A).

**Figure 1:**
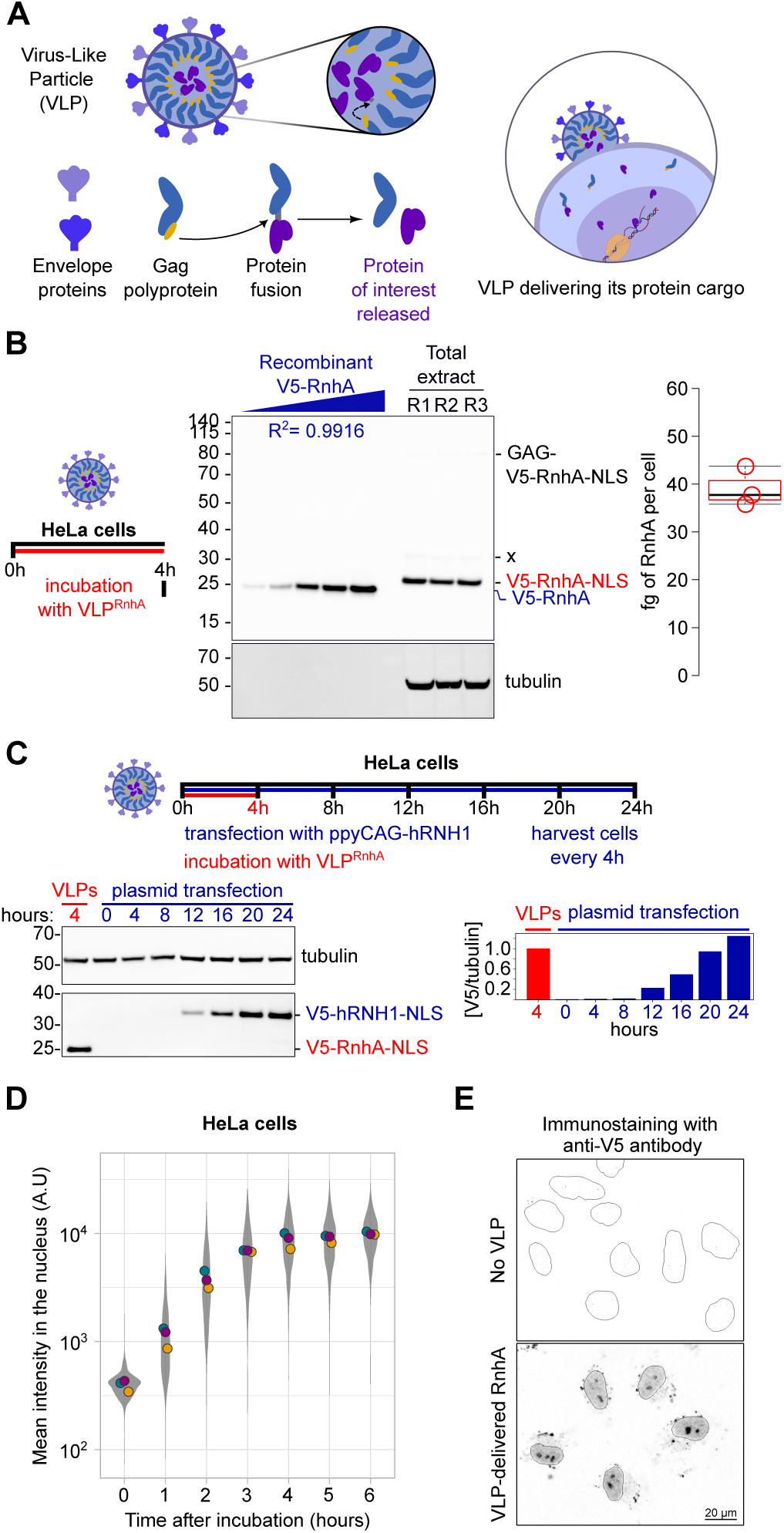
Virus-Like Particles (VLPs) rapidly deliver RnhA in the nuclei of live cells. All experiments were carried out in HeLa cells. **A.** Schematic representation of VLPs. The protease domain of the Gag polyprotein (in yellow) cleaves the junction between Gag and the protein of interest. As a result, VLPs deliver the ready-made protein of interest to target cells as a separate entity. **B**. Western blot analysis of VLP-delivered V5-RnhA-NLS after a 4-hour incubation. The first 5 lanes on the left correspond to increasing amounts of recombinant V5-RnhA protein lacking an NLS that were run as protein standards (R^2^ = 0.9916). The last three lanes correspond to total protein extracts from ∼1.5×10^5^ cells incubated with three independent lots of VLPs for 4 hours (R1-3). The position of the full-length GAG-RnhA fusion is indicated. The product marked ‘x’ corresponds to the product of a minority cleavage event within GAG rather than at the junction between GAG and RnhA. The fully cleaved product represents >95% of the total V5-tagged proteins in all three biological replicates. Tubulin was used as loading control. The estimated amount of VLP-delivered protein per cell is shown on the box and whisker plot on the right. **C.** Representative western blot analysis of the accumulation over time of V5-tagged, human RNase H1 (V5-hRNH1-NLS) upon transfection of an over-expression plasmid in cells, as indicated on the timeline above. The amount of V5-tagged RnhA (V5-RnhA-NLS) delivered by VLPs after four hours is shown for comparison. Tubulin was used as a loading control. V5/tubulin ratios (in blue) normalised over the VLP condition (in red) are shown on the right. **D.** Immuno-staining was used to quantify the nuclear levels of VLP-delivered RnhA over time in cells. The distribution of intensities in individual nuclei is shown in grey. The circles of different colours indicate the average nuclear intensity in 3 independent experiments. **E.** Representative images of the immuno-localisation of VLP-delivered RnhA at 4h. Cells not incubated with VLPs were used as controls. Nuclei borders are indicated by a solid line.

We produced VLPs carrying the RNase H1 enzyme from *Escherichia coli* (RnhA) fused at the N-terminus to the V5 epitope and at the C-terminus to a Nuclear Localisation Signal (NLS). The choice of RnhA was motivated by the fact that it has a high catalytic activity and that it is unlikely to be targeted by inactivating post-translational modifications or to interact with endogenous protein partners. RnhA was previously shown to recognise RNA:DNA hybrids *in situ* in permeabilised human cells (MapR procedure, (Yan *et al*, 2019)) or to remove RNA:DNA hybrids at tRNA genes in *Schizosaccharamyces pombe* (Legros *et al*, 2014). These results provide strong evidence that RnhA can recognise and process RNA:DNA hybrids in the chromatin of eukaryotic cells. Interestingly, the amount of RnhA loaded in VLPs can be adjusted during production for dose-dependent delivery and two proteins can be loaded and delivered at the same time and at different ratios, establishing VLPs as a flexible protein delivery system (Fig EV1A).

Western blot of total protein extracts confirmed that a 4-hour incubation with VLPs delivered a consistent amount of RnhA in live HeLa cells (Fig 1B). As a comparison, it was necessary to wait ∼20 hours to obtain a similar amount of ectopically expressed human RNase H1 (hRNH1) after plasmid transfection (Fig 1C). More than 95% of the VLP-delivered RnhA was cleaved off the Gag moiety, establishing that the majority of the additional RNase H activity in cells is provided by RnhA and not by the Gag-RnhA fusion (Fig 1B). To estimate the amount of RnhA delivered into HeLa cells after 4 hours, we used increasing amounts of recombinant V5-tagged RnhA as standards. Using linear regression, we could estimate that VLPs deliver ∼40 fg of RnhA per cell in 4 hours, which corresponds to ∼10^6^ molecules (Fig. 1B). As a comparison, it was previously estimated that there are only ∼200 and ∼5000 molecules of endogenous RNase H1 in resting and activated human T lymphocytes respectively (Wolf *et al*, 2020).

Immunostaining showed that VLP-delivered RnhA accumulated exponentially in the nuclei of HeLa cells in the first three hours of incubation before reaching a plateau after four hours (Fig 1D). In nuclei, RnhA accumulated in the nucleolus and was also uniformly distributed over chromatin, in a manner reminiscent of the endogenous RNase H1 enzyme (Shen *et al*, 2017) (Fig 1E). Fractionation of protein extracts established that ∼60% of VLP-delivered RnhA remained in the soluble fraction whilst ∼40% associated in a salt-sensitive manner with chromatin (Fig EV1B). Taken together, these data show that VLPs reproducibly deliver a ∼1000-fold excess of RNase H1 enzyme in live HeLa cells under 4 hours. We call this strategy ‘*iCoRD’* for ‘In <u>C</u>ellul<u>o</u> <u>R</u>Nase H <u>D</u>elivery’.

### *iCoRD*-delivered RnhA has a negligible impact on the steady state transcriptome

To evaluate the impact of *iCoRD*-delivered RnhA on the transcriptome, we carried out total RNA-seq analysis of HeLa cells treated with RnhA-containing VLPs for 4h or 24h (Fig EV2A). As controls, we used VLPs carrying mCherry or the inactive RnhA^D10N^ mutant (thereafter called *d*RnhA) that can recognise but not degrade RNA:DNA hybrids (Kanaya *et al*, 1990). We only detected a maximum of two transcripts whose levels were mildly altered upon addition of VLPs delivering either control proteins or active RnhA, even after a 24-hour incubation (Fig EV2BC). These observations were confirmed when sequencing chromatin-associated RNAs (Fig EV2D). In sharp contrast, re-analysis of published RNA-seq data (Tan-Wong *et al*, 2019) showed comprehensive transcriptome perturbations upon plasmid-driven expression of RNase H1 for 36 hours (Fig EV2E). In particular, transcription of the ubiquitin-like *ISG15*, a known regulator of DNA replication stress (Raso *et al*, 2020; Moro *et al*, 2023; Wardlaw & Petrini, 2022), was strongly activated in those cells (Fig EV2F). To conclude, contrary to the lengthy over-expression of RNase H1, there are no significant confounding effects on the steady state transcriptome associated with the VLP-mediated delivery of either control proteins or active RnhA.

### *iCoRD*-delivered *d*RnhA associates with RNA:DNA hybrids in cells

To determine whether *iCoRD*-delivered RnhA can recognise and associate with RDHs in human cells, we first tested whether it would co-localise with synthetic R-loops transfected into live cells (Fig 2A). To this end, we used a previously published strategy to produce R-loops at *mAirn* using *in vitro* transcription (IVT) (Carrasco-Salas *et al*, 2019). When transfected into HeLa cells, their detection by the S9.6 antibody yielded strong nuclear and cytosolic foci, also visible in the DAPI channel (Fig 2A). We first tested whether *d*RnhA or the W90A mutant (RnhA^W90A^), which was previously suggested to be defective for RNA:DNA hybrid binding (Kanaya *et al*, 1991), would co-localise with these synthetic R-loops *in situ*. As RnhA^W90A^ was less efficiently incorporated into VLPs than RnhA (Figure EV3), we first carried out this first experiment using a classic over-expression strategy. Both *d*RnhA and RnhA^W90A^ proteins were over-expressed at similar levels (Fig 2B). We found that 95% of S9.6 foci in cells co-localised with *d*RnhA (812 foci counted in 113 cells), whilst only 14% were stained with RnhA^W90A^, albeit weakly (787 foci counted in 104 cells) (Fig 2C). S9.6 co-localisation was also observed when *d*RnhA was delivered with VLPs, demonstrating that its inclusion into VLPs does not alter its ability to recognise RDHs (Fig 2D). These observations establish that *d*RnhA is able to recognise and associate with synthetic RNA:DNA hybrids in live cells.

**Figure 2:**
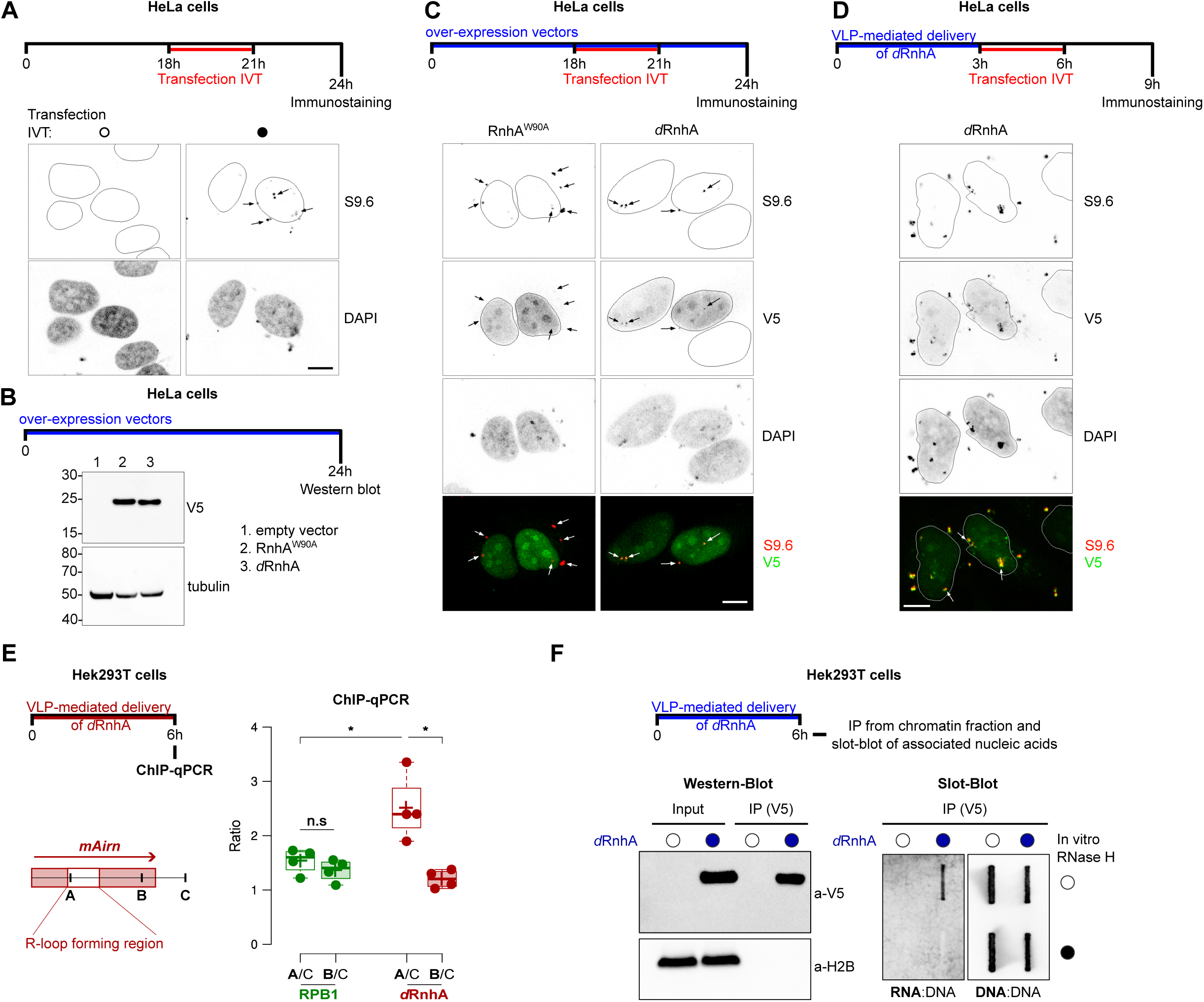
*iCoRD*-delivered *d*RnhA associates with RNA:DNA hybrids in cells. The cell lines used and the timeline of the experiment is indicated above each panel. **A.** The transfection of *in vitro* prepared R-loops results in the appearance of strong S9.6 foci (arrows). Nuclei borders are indicated by a solid line. Scale bar 10 µm. **B.** Western blot was used to confirm the similar levels of the indicated proteins in cells. Tubulin was used as loading control. **C.** Representative images of the co-localisation between the synthetic R-loops marked with the S9.6 antibody and the indicated V5-tagged proteins. Nuclei borders are indicated by a solid line. Scale bar 10 µm. **D.** Representative image of the co-localisation between the synthetic R-loops marked with the S9.6 antibody and *iCoRD*-delivered, V5-tagged *d*RnhA. Nuclei borders are indicated by a solid line. Scale bar 10 µm. **E.** ChIP-qPCR of the endogenous RPB1 or *iCoRD*-delivered V5-tagged *d*RnhA at *mAirn* expressed from an episome. The positions of the primers A, B and C used in the qPCR is indicated (n = 4). **F.** Chromatin-associated, VLP-delivered *d*RnhA was immuno-precipitated (left) and its associated nuclei acids analysed by slot blot (right).

We previously identified the region of *mAirn* that is most likely to form R-loops when transcribed *in vitro* (Carrasco-Salas *et al*, 2019). To strengthen the idea that *iCoRD*-delivered *d*RnhA could recognise RDHs *in situ*, we investigated whether *d*RnhA would be particularly enriched in this region. For this, we used a previously-published episome that expresses *mAirn* from a Doxycyclin (DOX)-inducible promoter (Hamperl *et al*, 2017) and monitored the enrichment of *iCoRD*-delivered *d*RnhA along the *mAirn* gene upon induction using Chromatin Immunoprecipitation followed by quantitative PCR (ChIP-qPCR). We used the endogenous Rpb1 subunit of RNA polymerase 2 (RNAP2) as control. While Rpb1 displayed a homogenous enrichment at all loci tested within *mAirn*, *d*RnhA accumulated significantly on the R-loop forming region (Fig 2E), strengthening the idea that *d*RnhA is able to recognise RDHs in live human cells.

### *iCoRD*-delivered *d*RnhA associates with RNA:DNA hybrids on chromatin

To determine whether *iCoRD*-delivered *d*RnhA could interact with endogenous, chromatin-associated RDHs, we first immunoprecipitated *d*RnhA from the chromatin-containing fraction and analysed the associated nucleic acids by slot blot for the presence of dsDNA and RDHs. RDHs were specifically enriched upon immunoprecipitation of *d*RnhA, confirming that VLP-delivered *d*RnhA associates with RNA:DNA hybrids in human cell extracts (Fig 2F).To determine whether *d*RnhA recognises the same RDHs as its human counterpart, we used ChIP-qPCR to measure the enrichment of *d*RnhA at loci displaying weak, medium and high accumulation of human RNase H1, as determined from published datasets (Chen *et al*, 2017) (Fig 3A). We confirmed the accumulation of VLP-delivered *d*RnhA but not mCherry or RnhA at these sites (Fig 3AB). Similarly, the hybrid binding-defective RnhA^W90A^ mutant failed to yield significant ChIP signals (Fig 3C). Taken together these observations indicate that ChIP signals of VLP-delivered *d*RnhA are highly likely to reflect the presence of genuine RNA:DNA hybrids.

**Figure 3:**
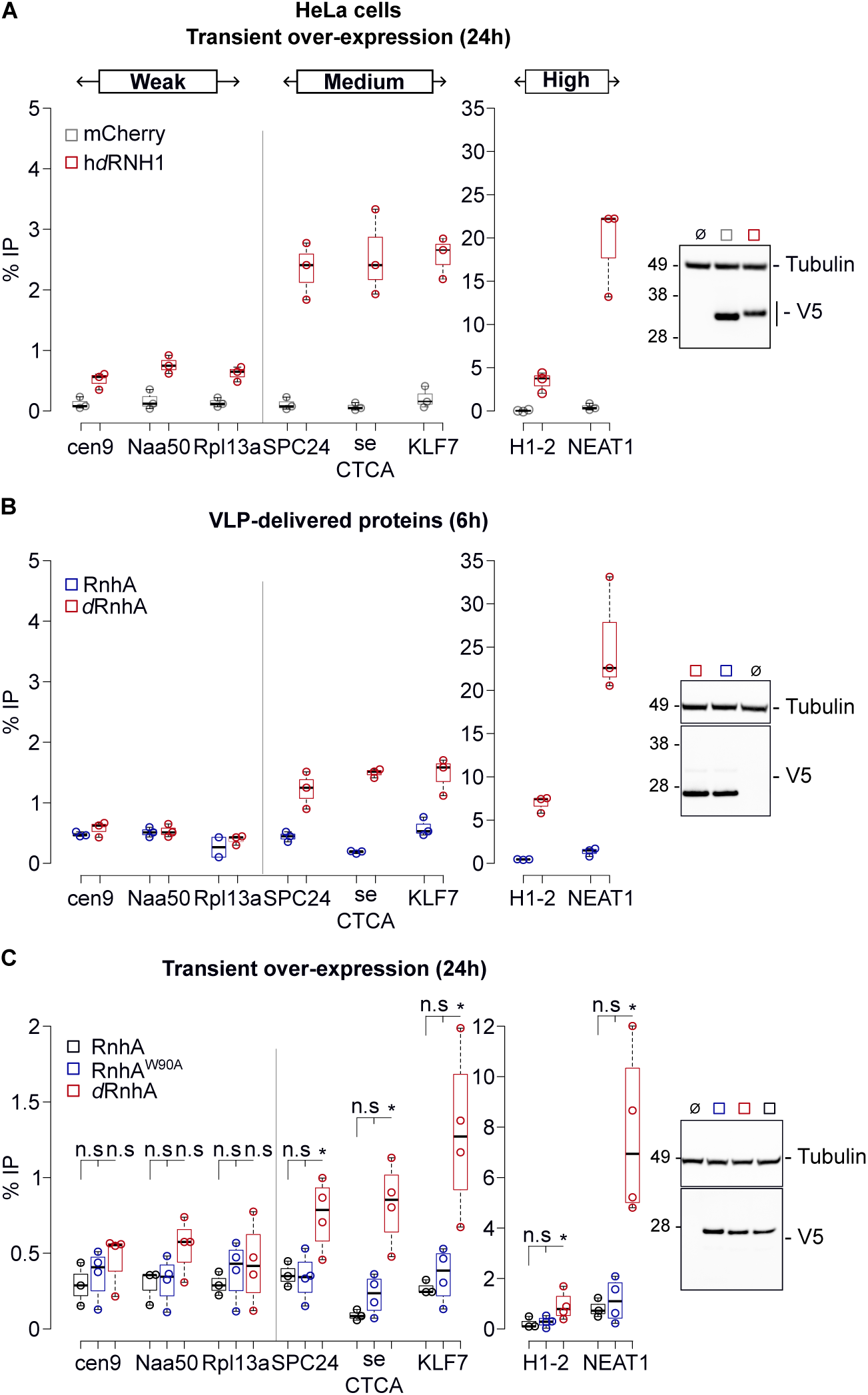
*iCoRD*-delivered *d*RnhA associates with endogenous RNA:DNA hybrids on chromatin. HeLa cells were used in all experiments. ChIP-qPCR analysis was performed for the indicated proteins. The loci analysed were selected because of their varying enrichment for the human RNase H1 (weak, medium and high). For each experiment, western blot was used to confirm the similar cellular levels of the different proteins analysed. Tubulin was used as loading control. **A.** ChIP-qPCR of V5-tagged hRNH1^D210N^ (h*d*RNH1) and V5-tagged mCherry, transiently over-expressed for 24h (n = 3). **B.** ChIP-qPCR of the indicated VLP-delivered proteins (6-hour incubation) (n = 3). **C**. ChIP-qPCR of V5-tagged RnhA, RnhA^W90A^ or *d*RnhA transiently over-expressed for 24h (n ≥ 3). For each locus, the Wilcoxon Mann-Whitney test was used to compare the distributions of values for RnhA^W90A^ or *d*RnhA with the distribution of values for RnhA. n.s = not significant; *: p-value ≤ 0.05.

To conclude, the microscopy and molecular assays presented in Figures 1–3 show that VLPs rapidly deliver different RnhA variants in the nuclei of live human cells, where they recognise and associate with both synthetic and endogenous RNA:DNA hybrids.

### *iCoRD*-delivered RnhA localises in the vicinity of active replication forks

As the over-expression of RNase H1 has been shown to modulate replication fork progression in response to replication stress, we tested whether VLP-delivered RnhA could be found in the vicinity of replication forks. We synchronised HeLa cells in S-phase using a single thymidine block (Fig 4A) and used Proximity Ligation Assay (PLA) to measure the proximity between VLP-delivered proteins and the replisome component Proliferating Cell Nuclear Antigen (PCNA). As the different VLP-delivered proteins carry the same V5 tag in their N-terminus and the same NLS in their C-terminus, PLA signals for the different VLP-delivered proteins are directly comparable. We first confirmed that the delivery of the different proteins did not impact the quality of the synchronisation in S phase (Fig 4B). Strikingly, only the delivery of the inactive *d*RnhA yielded strong PLA signals with PCNA (Fig 4C). To quantify PLA signals in an unbiased manner, we used a machine-learning approach to classify nuclei into four different clusters depending on the strength of their signals (see Methods) (Fig 4D). This confirmed that *d*RnhA yielded stronger PLA signals than RnhA (Fig 4E), despite immuno-staining experiments finding no major differences in the levels of RnhA or *d*RnhA on chromatin after the pre-extraction protocol used for the PLA (Fig 4F, left panel). In contrast, the levels of mCherry decreased strongly upon pre-extraction, as expected for a nucleosoluble protein (Fig 4F, left panel). The delivery of either mCherry, RnhA or *d*RnhA did not affect the levels of PCNA on chromatin (Fig 4F, right panel). This shows that the failure by RnhA to yield robust PLA signals cannot be explained by its depletion from chromatin. As *d*RnhA is unlikely to harbour determinants for targeting to specific genomic regions in human cells, its proximity to PCNA most likely reflects an association with fork-proximal RNA:DNA hybrids.

**Figure 4:**
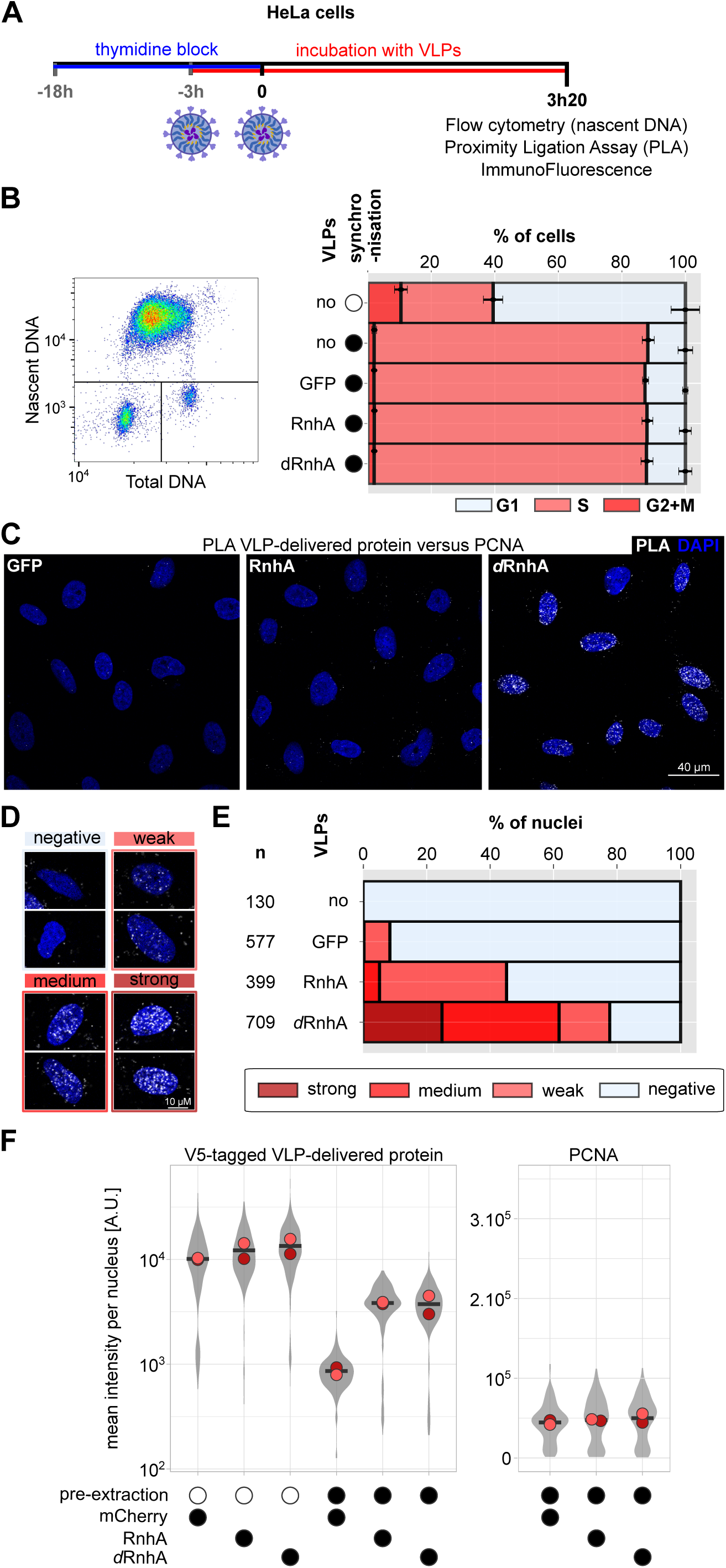
*iCoRD*-delivered RnhA is positioned in the vicinity of active replication forks. All experiments were carried out in HeLa cells. **A.** Schematic representation of the experiment. **B.** Cell-cycle profile of cells synchronised in S-phase and incubated with VLPs delivering the indicated proteins. (left) example of a typical FACS profile. (right) % of cells in the different cell-cycle phases (n≥3). Error bars correspond to the standard deviation. **C.** Examples of PLA signals in the indicated conditions. **D.** A machine learning approach was used to sort nuclei into different clusters of varying PLA signals. Representative nuclei for the different clusters. **E.** % of nuclei in the different clusters per conditions. The total number of nuclei analysed over 3 to 5 biological replicates is indicated on the left (n) **F**. Immunostaining was used to quantify the nuclear levels of VLP-delivered RnhA and *d*RnhA before and after pre-extraction. mCherry was used as a control for a soluble protein sensitive to the pre-extraction. PCNA levels on chromatin were also measured in the different conditions (right). The distribution of intensities in individual nuclei is shown in grey. The circles of different colours indicate the average nuclear intensity in 2 independent experiments.

### *iCoRD*-delivered RnhA rescues the progression of replication forks under stress

Having established that *iCoRD*-delivered RnhA is enriched in the vicinity of replication forks, we asked whether it can rescue the progression of replication forks in response to stress. We first incubated asynchronous U2OS cells with VLPs carrying either mCherry, RnhA or *d*RnhA. After 3 hours of incubation, cells were subjected to consecutive pulses with the thymidine analogues IdU and CldU (Fig EV4). During the CldU pulse, cells were exposed or not to 50 µM Hydroxyurea (HU) to induce fork slowdown (Andrs *et al*, 2023; Somyajit *et al*, 2017). As expected, in cells pre-treated with VLPs carrying the control protein mCherry, CldU tracks were significantly shorter than IdU tracks in the presence of HU (CldU/IdU ratio ∼ 0.65) (Fig 5A). Strikingly, cells pre-treated with RnhA-containing VLPs were completely resistant to this dose of HU (CldU/IdU ratio ∼ 1). This rescue required the catalytic activity of RnhA because cells pre-treated with *d*RnhA-containing VLPs behaved like cells treated with mCherry-containing VLPs (Fig 5A). The rescue effect of the over-expression of human RNase H1 on replication fork progression in response to 50 µM HU was shown to require Primpol activity (Andrs *et al*, 2023). Here we show that Primpol is also required for the HU-resistant progression of replication forks in U2OS cells treated with RnhA-containing VLPs (Fig EV5A). Taken together, these observations confirm that *iCoRD*-delivered RnhA is active in cells and promotes the Primpol-dependent progression of replication forks in response to low doses of HU in the same way as human RNase H1. Using a similar strategy, we also show that RnhA-containing VLPs can rescue the replication fork slowdown associated with loss of Topoisomerase I (Top1) in HeLa cells (Tuduri *et al*, 2009; Promonet *et al*, 2020) (Fig EV5B).

**Figure 5:**
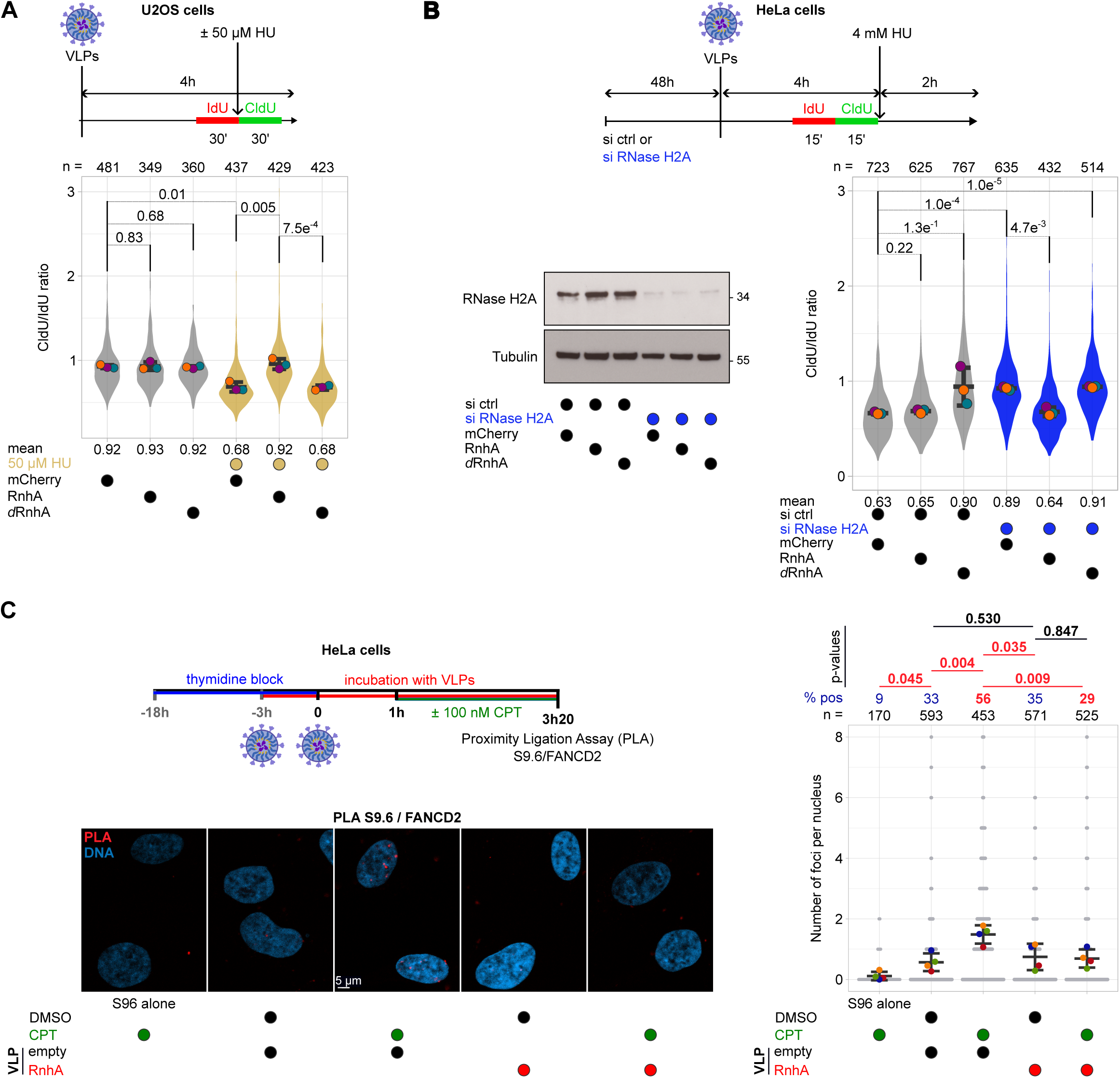
*iCoRD*-delivered RnhA rescues the progression and stability of replication forks under stress. The cell line used and the timeline of the experiment is shown for each panel. (**AB**) After short pulses of the indicated duration with IdU and CldU, chromatin spreading was used to monitor replication fork progression (see also Fig EV4). The ratio CldU/IdU for individual fibres is represented on superplots, as recommended previously (Lord *et al*, 2020). The distribution of values is shown in grey and the circles of different colours indicate the mean values in 3 independent experiments. Comparisons between the means of the replicates were carried out and Welch’s t-test was performed to calculate the indicated p-values. In (**B**), western blot was used to confirm the depletion of RNase H2A. Tubulin was used as loading control. The impact of VLPs delivering the indicated proteins on the CldU/IdU ratio was assessed in **(A)** in cells treated with 50 µM HU for 30 mins and in **(B)** in cells depleted or not of RNase H2A and treated with 4 mM HU. The total number of chromatin fibres that were scored is indicated at the top of graphs. **(C)** Results of the PLA between FANCD2 and S9.6 (n =4); (left) representative nuclei in all conditions; (right) number of PLA foci per nucleus. The total number of nuclei scored and the % of cells displaying PLA foci are also indicated at the top of the graph. Comparisons between the means of the replicates were carried out and Welch’s t-test was performed to calculate the indicated p-values.

We previously demonstrated that loss of RNase H2A interferes with the resection of stalled replication forks in response to high doses of HU (Heuzé *et al*, 2023). We show here that *iCoRD*-delivered RnhA could also rescue this defect (Fig 5B). Interestingly, the delivery of the inactive *d*RnhA mutant had a dominant-negative effect on fork resection, mimicking to some extent loss of RNase H2A (Fig 5B). These observations are consistent with the idea that active RnhA could remove resection-impairing post-replicative RNA:DNA hybrids, whilst the inactive *d*RnhA might stabilise them. This model is in agreement with the fact that PLA signals with PCNA were stronger for *d*RnhA than for RnhA (Fig 4). Taken together, these results demonstrate that *iCoRD*-delivered RnhA is active in human cells, that it can compensate for the loss of RNase H2A and that it modulates in *-cis* the behaviour of replication forks in response to a diverse array of replication stresses.

### *iCoRD*-delivered RnhA degrades fork-proximal RNA:DNA hybrids

The above data suggest that *iCoRD*-delivered RnhA degrades in *-cis* the fork-proximal RNA:DNA hybrids that impede the continuous progression of replication forks upon stress. To strengthen this observation by a different approach, we used Proximity Ligation Assay (PLA) to monitor the proximity between FANCD2, a marker of stressed replication forks (Lossaint *et al*, 2013), and S9.6, an antibody that recognises RNA:DNA hybrids (Boguslawski *et al*, 1986). HeLa cells were synchronised in S phase and incubated with *iCoRD* as above (Fig 5C). Replication stress was then induced or not by treating cells with 100 nM of the Topoisomerase I poison Camptothecin (CPT), which was shown previously to induce the formation of both FANCD2 foci and RNA:DNA hybrids (Chappidi *et al*, 2020). As expected, CPT treatment triggered a small, albeit reproducible and significant increase both in the number of cells exhibiting PLA foci (% pos, Fig 5C) and in the number of PLA foci per nucleus (Fig 5C). Consistent with our hypothesis, both increases were fully reversible with *iCoRD*-delivered RnhA (Fig 5C). These data are consistent with the idea that *iCoRD*-delivered RnhA can degrade FANCD2-proximal RDHs upon CPT treatment.

### *iCoRD*-delivered RnhA has no significant impact on R-loops mapped by DRIP or Cut&Tag

It is commonly assumed that the over-expression of RNase H1 degrades the cotranscriptional R-loops that impede fork passage. As the S9.6-based DNA:RNA Immuno-precipitation (DRIP) method is frequently used to map R-loops, we tested whether *iCoRD*-delivered RnhA could also remove DRIP signals. We first used the S9.6 antibody to measure the global amount of RNA:DNA hybrids in genomic DNA (gDNA) blotted on a membrane. To assess whether RnhA could remove excessive R-loops, R-loop formation was strongly induced by a short treatment with high doses of CPT, as described previously (Saha *et al*, 2022). Surprisingly, S9.6 signals were largely unaffected by the delivery of RnhA, whether or not cells were treated with CPT, whilst they were fully sensitive to an *in vitro* RNase H treatment (Fig EV6A). Similarly, DNA:RNA IP (DRIP-qPCR) signals at canonical R-loop-forming loci were sensitive to an *in vitro* treatment with RNase H but resistant to the delivery of RnhA by VLPs (Fig EV6B).

We sought to confirm these unexpected observations using high-resolution, strand-specific and calibrated DRIP-seq in cycling U2OS cells (Fig 6A-C). As expected, DRIP signals were concentrated around Transcription Start Sites (TSS) and Transcription End Sites (TES) on both DNA strands (Fig 6BC) and were fully sensitive to an *in vitro* treatment with commercial RNase H (Fig 6A). However, we found no difference in DRIP signals when cells were incubated with GFP- or RnhA-carrying VLPs (Fig 6D), confirming our DRIP-qPCR results at the genome-wide scale (Fig EV6B). Similarly, we found that S9.6 Cut&Tag signals were indistinguishable in cells incubated with mCherry- or RnhA-carrying VLPs (Fig EV6CD).

**Figure 6:**
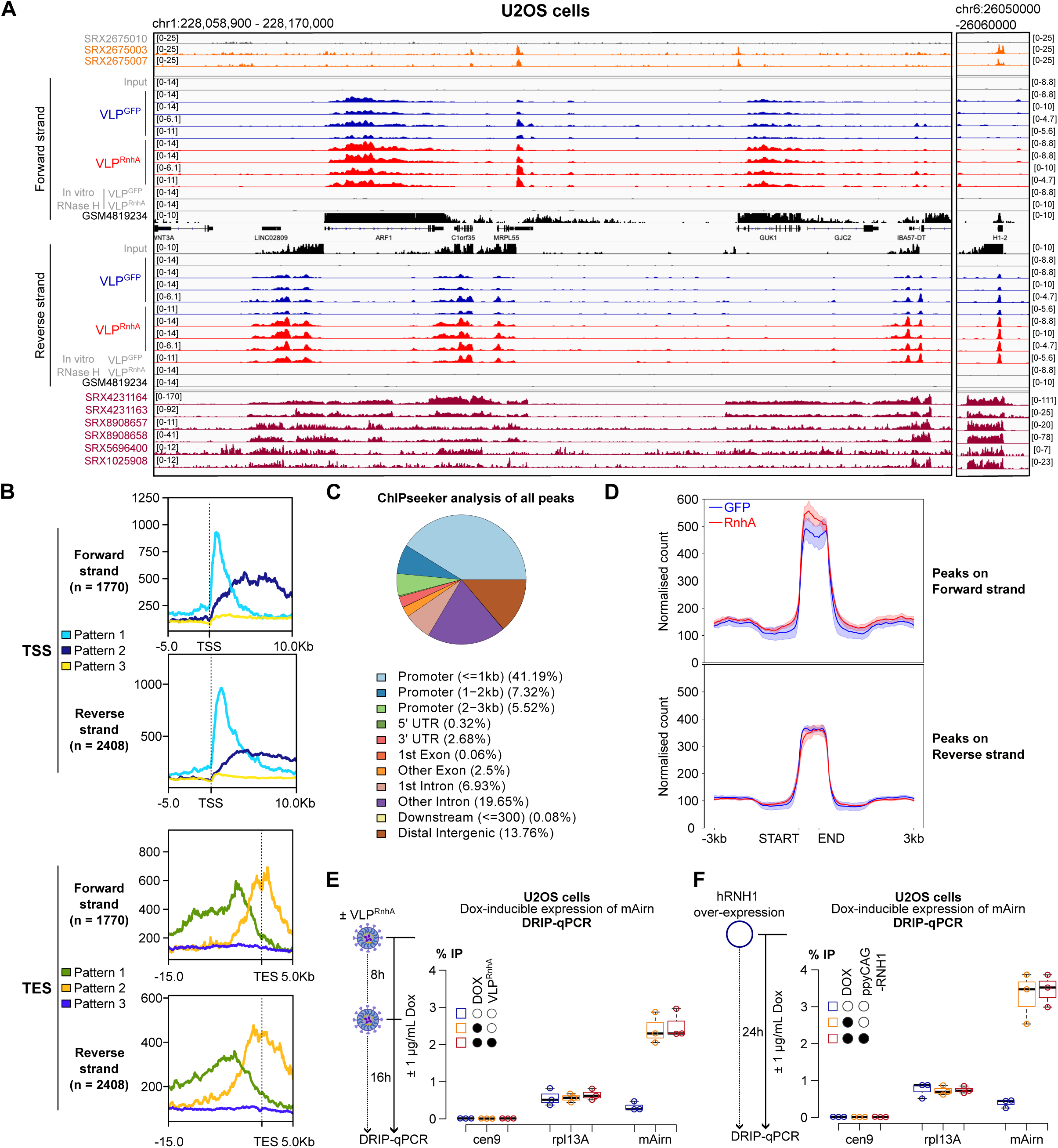
The significant increase in nuclear RNase H1 levels does not impact DRIP signals. **A.** Examples of cpm-normalised and strand-specific DRIP-seq signals upon treatment with GFP-containing or RnhA-containing VLPs (blue and red tracks respectively). Input samples and DRIP samples treated *in vitro* with commercial RNase H are shown as controls (grey tracks). For comparison, published R-ChIP tracks from HEK293T cells (SRX2675010 = negative control, SRX2675003 & SRX2675007 = R-ChIP (Chen *et al*, 2017)) and published DRIP-seq tracks from U2OS cells (SRX4231163 & SRX4231164, (De Magis *et al*, 2019); SRX8908657 & SRX8908658, (Villarreal *et al*, 2020); SRX5696400, (Wu *et al*, 2020); SRX1025908, (Gorthi *et al*, 2018)) are also shown. TT-seq tracks (GSM4819234, (Sawicka *et al*, 2021)) show nascent transcription at the loci of interest in U2OS cells. **B.** Metaplots of DRIP signals around TSS or TES for DRIP-positive genes on the forward or reverse DNA strand. k-means clustering was used to group genes displaying similar distribution patterns of DRIP signals (3 patterns for each TSS or TES location). Similar distribution patterns were identified on both DNA strands. **C.** ChIPseeker analysis of the distribution of DRIP signals with respect to the indicated features (all peaks). **D.** Calibrated DRIP signals around called peaks (see Methods). **EF.** U2OS-*mAirn* cells were treated for 24 hours with doxycycline to induce the expression of *mAirn*. qPCR was used to quantify DRIP signals at *mAirn* and at two control loci (cen9 as negative control and Rpl13A as positive control) either (**E**) upon VLP-mediated delivery of RnhA over a 24-hour period or (**F**) upon the over-expression of human RNase H1 for 24 hours.

To test whether VLP-delivered RnhA could remove DRIP signals induced by the *de novo* transcription of an R-loop forming gene, we used a cell line in which five copies of the R-loop forming *mAirn* gene were integrated under the control of a Dox-inducible promoter (Werner *et al*, 2025). The addition of Dox for 24 hours induced a reproducible ∼10-fold increase in DRIP signals at *mAirn* but this increase also occurred when RnhA-containing VLPs were present during the whole duration of induction (Fig 6E). This shows that *iCoRD*-delivered RnhA could not antagonise the increase in DRIP signals associated with *mAirn* transcription, even after a 24-hour treatment.

We wondered whether the classic over-expression of human RNase H1 (hRNH1) would be able to reduce DRIP signals. The 24-hour over-expression of hRNH1 had no impact on the S9.6 signals measured by slot blot, even after R-loop formation was strongly induced by a short treatment with high doses of CPT (Fig EV7AB). These observations were confirmed by DRIP-qPCR (Fig EV7C). Similarly, the 24-hour over-expression of hRNH1 did not antagonise the increase in DRIP signals associated with the induction of *mAIRN* transcription (Fig 6F, Fig EV7D).

Taken together, our data demonstrate that high levels of active RNase H1 in live nuclei do not affect scheduled or unscheduled RNA :DNA hybrid levels, as measured by slot blots, DRIP or Cut&Tag.

### *iCoRD*-delivered RnhA removes *d*RnhA R-ChIP signals

As our data showed clearly that *iCoRD*-delivered RnhA can rescue fork progression under stress (Fig 5AB and Fig EV5) and recognise and degrade RNA:DNA hybrids in normal (Fig 2&3) or stressed conditions (Fig 5C), we were surprised that it had no impact on DRIP signals, whether scheduled or unscheduled. We therefore sought to confirm this result by investigating whether a large excess of *iCoRD*-delivered RnhA could eliminate *d*RnhA R-ChIP signals but not DRIP signals at the same loci. To test whether *iCoRD*-delivered RnhA could eliminate *d*RnhA R-ChIP signals, we incubated cells in the presence of 2 different combinations of VLPs: either (i) VLPs delivering untagged mCherry together with VLP delivering V5-tagged *d*RnhA at a ratio [10:1] or (ii) VLPs delivering Flag-tagged RnhA together with VLP delivering V5-tagged *d*RnhA at a ratio [10:1] (Fig 7A). Strikingly, *d*RnhA ChIP signals at high accumulation sites (Fig 3) were sensitive to a 10-fold excess of active RnhA (Fig 7A). Considering that RnhA does not stably associate with these loci (Fig 3B), this observation strongly indicates that the excess of RnhA degrades the RNA:DNA hybrids that recruit *d*RnhA. Importantly, at the same sites, the same amount of VLP-delivered RnhA had no impact on DRIP signals (Fig 7B). These observations confirm that VLP-delivered RnhA is active against RDHs on chromatin, but has no detectable impact on DRIP signals.

**Figure 7:**
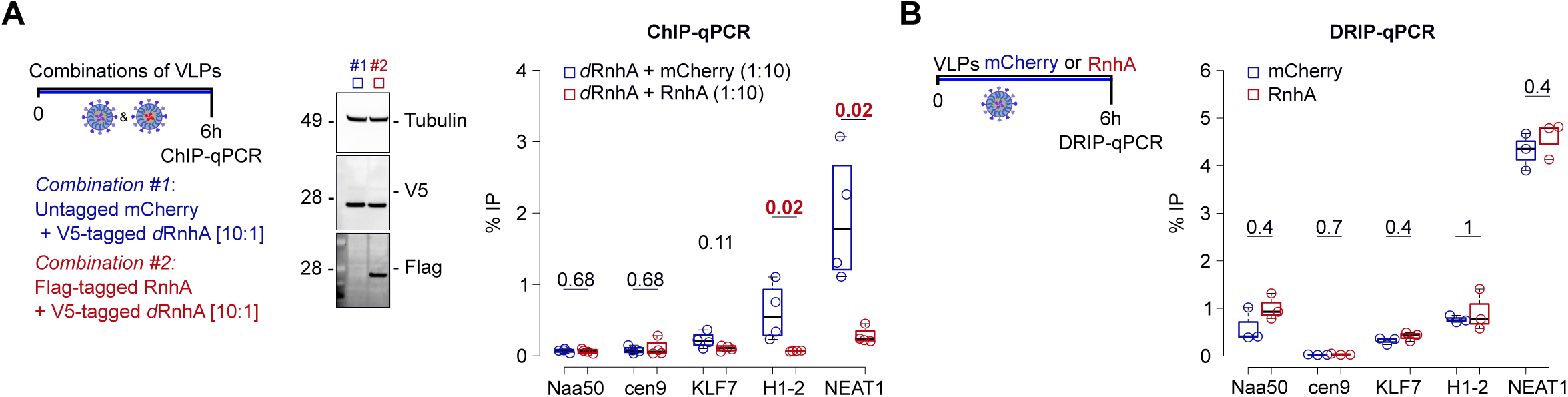
At *d*RnhA-rich loci, *d*RnhA R-ChIP but not DRIP signals are sensitive to an excess of RnhA. All experiments were carried out in HeLa cells. **A.** Cells were incubated for 6 hours with two different combinations of VLPs at a ratio of [10:1]: either untagged mCherry and V5-tagged *d*RnhA (blue) or Flag-tagged RnhA and V5-tagged *d*RnhA (red). ChIP-qPCR was then performed with V5-tagged *d*RnhA (n = 4). For each locus, the Wilcoxon Mann-Whitney test was used to compare the distributions of values. The resulting p-values are indicated on the graph. **B.** DRIP was carried out at the same loci after a 6-hour delivery of either VLP^mCherry^ or VLP^RnhA^ (n = 3). For each locus, the Wilcoxon Mann-Whitney test was used to compare the distributions of values. The resulting p-values are indicated on the graph.

## DISCUSSION

We set up *iCoRD* to engineer the rapid and quantitative delivery of RnhA variants in live cells. We demonstrated that *iCoRD*-delivered RnhA can degrade the population of RNA:DNA hybrids that normally impede fork resection and restart upon replication stress but not the population of hybrids that gives rise to DRIP or R-loop Cut&Tag signals, even in stressed conditions. We also found that DRIP signals were largely resistant to the prolonged over-expression of human RNase H1. Our work therefore establishes that DRIP signals are sensitive to RNase H1 *in vitro* but largely resistant to a large excess of RNase H1 *in situ*. We conclude from these observations that RNA:DNA hybrids mapped by DRIP are likely to correspond to a separate class that is unlikely to contribute to the genomic instability that can be corrected by increasing RNase H1 activity in cells.

### *iCoRD*, a promising method to study RNase H1-sensitive RNA:DNA hybrids in cells

We have shown that Virus-Like Particles (VLPs) can be used to deliver controllable amounts of ready-made sensors (*d*RnhA) and regulators (RnhA) of RNA:DNA hybrids in live human cells, with both unprecedented speed and efficiency and negligible effects on the steady state transcriptome. Importantly, no purification of recombinant protein is necessary for this approach and multiple proteins can be delivered at the same time (Fig EV1A). This methodology does not require any genetic modification of target cells and the delivery in parallel of a similar amount of control proteins (GFP, mCherry) is easily carried out. This makes *iCoRD* particularly attractive to study the role of RNA:DNA hybrids in primary cells.

Here we chose to deliver non-human proteins to limit the risk that the RNA:DNA sensors and regulators of interest could be inactivated by post-translational modifications or that their distribution or function could be impacted by interactions with endogenous proteins. For example, it was shown previously that RPA associates with and stimulates the activity of human RNase H1 but not *E. coli* RnhA (Nguyen *et al*, 2017). This means that a confounding effect of over-expressing human RNase H1 could be to titrate RPA. This risk is avoided with the *iCoRD*-mediated delivery of RnhA.

### *iCoRD* demonstrates the direct impact of RNA:DNA hybrids on fork transactions

As reported both previously (Tan-Wong *et al*, 2019; Chao & Fazzio, 2026) and here (Fig EV2E), the prolonged over-expression of RNase H1 could have a significant impact on the steady-state transcriptome in metazoans. It is difficult to evaluate how much those transcriptome alterations could indirectly contribute to the rescue of stressed replication forks by the prolonged over-expression of RNase H1. We note that the analysis of a previously published RNA-seq dataset (Fig EV2EF) revealed that the transcription of ISG15 was strongly activated upon prolonged RNase H1 over-expression in HeLa cells (Tan-Wong *et al*, 2019). As ISG15 conjugation to fork-associated proteins was shown to mitigate the consequences of replication stress (Raso *et al*, 2020; Moro *et al*, 2023; Wardlaw & Petrini, 2022), the sole activation of ISG15 might be sufficient to explain why RNase H1 over-expression alleviates the detrimental consequences of TRCs in those cells, independently of the hypothetical removal of RNA:DNA hybrids. This result therefore undermined the widely-accepted model that hybrids play a direct role at stressed replication forks. However, the observations that (i) that the short-term and transcriptome-neutral delivery of RnhA with *iCoRD* specifically at the time of DNA replication is sufficient to fully rescue the progression of replication forks upon replication stress (Fig 5) and (ii) that *iCoRD*-delivered RnhA is enriched at active replication forks (Fig 4), strongly suggest that RNA:DNA hybrids do indeed directly impact the processing of stressed replication forks in *-cis*.

### *iCoRD*-delivered RnhA manipulates RNA:DNA hybrids in the vicinity of stressed replication forks

Our functional assays have established that *iCoRD*-delivered RnhA exhibits RNase H activity in human cells. We further propose that *iCoRD*-delivered RnhA can act on at least two distinct, spatially separate populations of RNA:DNA hybrids depending on the stress condition and the cellular context. We suggested previously that in RNase H2A-depleted cells treated with 4 mM HU, the relevant hybrid population is most likely post-replicative and would form upon fork reversal (Heuzé *et al*, 2023). We have shown that *iCoRD*-delivered RnhA can substitute for the missing RNase H2 activity to degrade these RNA:DNA hybrids and restore normal resection of nascent DNA (Fig 5B). In contrast, catalytically-dead *d*RnhA might bind, but not resolve, these post-replicative hybrids, exerting a dominant-negative effect on fork resection (Fig 5B). In RNase H2-proficient cells exposed to 50 µM HU (Fig 5A and EV5A), we favor instead the model proposed by Andrs *et al*. (Andrs *et al*, 2023), in which RnhA/hRNH1 degrades a RNA:DNA hybrid formed ahead of the fork at a site of transcription-replication conflict. In this case, fork restart would require PrimPol-mediated repriming (Fig EV5A). Directly comparing the genomic location, structural context and genetic dependence of these different classes of RNA:DNA hybrids is an important question for future work.

### DRIP and Cut&Tag signals are resistant to high levels of RNase H1 in the nucleus

We have shown by microscopy and molecular assays that *iCoRD*-delivered *d*RnhA can recognise both synthetic and endogenous RNA:DNA hybrids in cells (Fig 2 and 3) and we provided evidence that *iCoRD*-delivered RnhA could degrade FANCD2-proximal RNA:DNA hybrids upon stress (Fig 5). However, our results indicate that a large excess of RnhA or hRNH1 in live nuclei could not suppress DRIP or Cut&Tag signals in cells, even when signals were increased, for example with high doses of CPT (Saha *et al*, 2022) or by the induction of *mAirn* (Fig. 6). Comparable observations were made recently by the Chédin and Crouch labs in very different experimental settings in both mice and human cells (preprint: Sakhuja *et al*, 2025). Similarly in yeast, the over-expression of RNase H1 had no impact on DRIP signals in unperturbed conditions and only a minor impact at specific tDNA genes in the absence of both RNase H1 and RNase H2 (Wagner *et al*, 2025).

It has long been known that the overlap between DRIP and RNase H1 ChIP signals is small (Chen *et al*, 2017), which could suggest that RNase H1 *in vivo* does not reliably target loci that otherwise exhibit strong DRIP signals, even when over-expressed. However, even at loci where both RnhA and hRNH1 were significantly enriched in ChIP experiments (Fig 3), RnhA failed to suppress DRIP signals *in situ* (Fig 7), suggesting that lack of accessibility is not the primary reason why DRIP signals are resistant to RNase H1 *in vivo*. Furthermore, neither RnhA nor hRNH1 could antagonise ‘nascent’ DRIP signals when induced *de novo* at the model R-loop forming *mAirn* gene or upon CPT treatment (Fig 6), ruling out that only ‘mature’ R-loops could be resistant to RNase H1 *in vivo*.

If limited accessibility of RNase H1 is not the primary explanation for these observations, we consider three alternative possibilities to explain why DRIP signals are sensitive to RNase H *in vitro but* resistant to RNase H1 *in situ*: either (i) the rate of R-loop formation may exceed the capacity of RNase H1 to remove them, even when the enzyme is present in large excess; (ii) the RNA moiety in R-loops might become rapidly resistant to RNase H1 *in vivo*, possibly because of a labile modification that would not withstand the DRIP procedure; or (iii) a substantial proportion of DRIP and Cut&Tag signals might not correspond to genuine R-loops. Early reports suggested that recombinogenic, transcription-dependent and RNA-containing structures called ‘R-loops’ could form artefactually in test tubes after DNA extraction from cells (Matsumoto & Ikeda, 1983). Although these structures were not shown to be sensitive to RNase H1 at the time, similar processes could underlie a significant fraction of current DRIP signals. Lack of cross-linking in the DRIP or Cut&Tag procedures might induce post-lysis R-loop formation or extension, which could mask the fraction of the signal that is genuinely sensitive to RNase H1 *in vivo* (see also (Belotserkovskii & Hanawalt, 2022)). We currently do not have strong arguments to favour either of these three hypotheses.

### *d*RnhA ChIP as an alternative to DRIP

Even if we do not yet have a biochemical explanation for these observations, they allow us to draw some important conclusions: (i) the fact that DRIP signals are sensitive to RNase H *in vitro* only rigorously proves that RNA:DNA hybrids were present in test tubes at the time of immuno-precipitation, but does not prove nor disprove that they were also present in cells; (ii) modulations of DRIP or Cut&Tag signals cannot explain an RNase H1-sensitive phenotype unless it is clearly demonstrated that the same signals are also significantly suppressed by increasing RNase H1 levels in cells.

The sensitivity of *d*RnhA ChIP signals to *iCoRD*-delivered RnhA (Fig 7) suggest that the transient delivery of *d*RnhA can be a convenient alternative to DRIP for mapping RNase H1-sensitive RNA:DNA hybrids on chromatin. Consistent with our previous observations in the yeast *Schizosaccharomyces pombe* (Legros *et al*, 2014), the active RnhA failed to yield significant ChIP signals. We speculate that it might interact too dynamically with its substrates to be quantitatively cross-linked to chromatin, possibly because it lacks a dedicated Hybrid-Binding Domain (HBD). As a result, it can serve as an effective negative control in R-ChIP approaches. Importantly, *iCoRD* reduces the risk of the artefactual stabilisation of RNA:DNA hybrids by the lengthy over-expression of an inactive RNase H1 mutant and is a promising alternative to current strategies to perform R-ChIP. As demonstrated on Figure 7A, it is easily tested using combinations of VLPs whether these *d*RnhA ChIP signals are sensitive to RNase H1 *in situ*.

## METHODS

### Cell culture and drug treatment

HEK293T, HeLa and U2OS cells were grown in DMEM medium supplemented with 10% (v/v) foetal calf serum and 1% (v/v) of Penicillin-Streptomycin (Gibco, cat. 15140-122) at 37°C with 5% CO_2_. The U2OS cells used to generate the DRIP-seq data were grown in McCoy’s 5A (Merck, cat. M9309) instead of DMEM medium. 1µg/mL puromycin (Gibco, cat. A11138-03) was added to the medium to amplify U2OS cells expressing *mAirn*, as described previously (Werner *et al*, 2025). 1µg/mL puromycin and 50 µg/mL hygromycin (Gibco, cat. 10687010) were added to the medium to amplify U2OS cells expressing GFP-RNase H1 as described previously (Andrs *et al*, 2023). (S)-(+)-Camptothecin (Merck, cat. C9911) was added at 20 µM final concentration in the growing medium and incubated for 5 minutes. DMSO (Invitrogen, cat. D12345) was used as control. Hydroxyurea (Merck, cat. H8627) was dissolved in the growing medium before being used at the final concentrations of either 50 µM (Fig 5A and EV5A) or 4 mM (Fig 5B). *mAirn* expression was induced with 1 µg/mL doxycycline (Sigma-Aldrich, cat. D3447) for 24 hours (Fig 6EF). GFP-RNase H1 expression was induced with 0.1 µg/mL doxycycline for 24 hours (Fig EV7), as described previously (Andrs *et al*, 2023).

### Protein expression and purification

the expression and purification of V5-tagged RnhA from *Escherichia coli* BL21(DE3) strain was carried out by ProteoGenix SAS (Schiltigheim, France). The concentration of the purified protein was carefully estimated on SDS-PAGE and Coomassie staining using decreasing amounts of BSA.

*Protein depletion:* Top1 and RNase H2A were depleted as previously described, in (Promonet *et al*, 2020) and in (Heuzé *et al*, 2023) respectively. Primpol was depleted using the siRNA GAGGAAACCGUUGUCCUCAGUGUAUUU (Horizon).

### RNH1 over-expression

∼4×10^5^ cells were transfected with 1µg of plasmid ppyCAG-hRNH1 (Addgene #111906) using jetOPTIMUS (Polyplus, cat. 101000025) according to the manufacturer’s instructions, and incubated for 24 hours. The over-expression of GFP-hRNH1 (Fig EV7) was carried out as described previously (Andrs *et al*, 2023). Both constructs allow the over-expression of the version of hRNH1 that lacks the mitochondrial localisation signal.

### Plasmid Construction

Plasmids encoding fusion proteins between Gag (MLV) and the protein of interest were derived from the BIC-Gag-Cas9 plasmid (Addgene plasmid #119942). To generate these constructs, the donor plasmid was digested with AgeI and KpnI to excise the Cas9 coding sequence. The resulting linearised plasmid backbone was purified via gel extraction. Inserts corresponding to the coding sequence of each protein of interest were amplified by PCR, incorporating homology arms complementary to the AgeI- and KpnI-digested vector termini. Assembly of the recombinant plasmids was performed using the NEBuilder HiFi DNA Assembly kit (NEB, cat. E5520S), following the manufacturer’s instructions. pBS-CMV-gagpol was a gift from Patrick Salmon (Addgene plasmid # 35614). pCMV-VSV-G was a gift from Bob Weinberg (Addgene plasmid # 8454) (Stewart *et al*, 2003). pBaEVRless was a gift from Els Verhoeyen (Girard-Gagnepain *et al*, 2014).

### VLP preparation and transduction

VLPs were prepared as described previously (Mangeot *et al*, 2019, 2021) with minor modifications. Plasmids were transfected into HEK293T cells plated at 3.5 × 10^6^ cells per 10 cm plate 24 h before transfection with the JetPrime reagent (Polyplus) according to the manufacturer’s instructions. 1 µg of the plasmid carrying the fusion between GagMLV and the protein of interest, 2.7 µg of pBS-CMV-gagpol, 0.3 µg of pCMV-VSV-G and 0.35 µg of pBaEVRless (Girard-Gagnepain *et al*, 2014) were co-transfected. Supernatants were collected after 40-48h and VLPs were purified and concentrated as previously described (Mangeot *et al*, 2019, 2021). Purified VLPs were incubated at 10 µL/mL in growing medium containing 10 µg/mL of polybrene (Merck, cat. TR-1003-G). For longer incubations, the growing medium was replaced with fresh growing medium containing new VLPs after 8 hours.

### Protein extracts and Western blot

Cell pellets were resuspended in 1X Sample buffer at a ratio of 4×10^6^ cells/mL. 1X sample buffer was prepared by diluting 4X Bolt LDS Sample Buffer (Invitrogen, cat. B0007), 10X Bolt LDS Sample Reducing Agent (Invitrogen, cat. B0009) and cOmplete, EDTA-free protease inhibitor cocktail tablets (Roche, cat. 05056489001) in ultrapure water. 20 µL of samples were run on Bolt 4-12 %, Bis-Tris Plus WedgeWell gels (Invitrogen, cat. NW04122BOX) and proteins were transferred on 0.2 µm nitrocellulose blotting membrane (Cytiva, cat. 10600015). Membranes were blocked with 5% milk diluted in PBS containing 0.02% Tween20. After incubation with the appropriate antibodies, signals were revealed using SuperSignal West Femto Maximum Sensitivity Substrate (ThermoScientific, cat. 34096). Fractionation of protein extracts was carried out using the Qproteome Nuclear Protein Kit (Qiagen, cat. 37582), according to the manufacturer’s instructions. The primary antibodies used were: SV5-Pk1 (BIO-RAD, cat. MCA1360), anti-Rrm2 (Proteintech, cat. 11661-1-AP), anti-histone H3 (Abcam, cat. ab1791), anti-GAG (Abcam, cat.130757), anti-SP1 (Proteintech, cat. 21962-1-AP). The anti-tubulin was purified from TAT-1 cells (RRID:CVCL_T980) (Woods *et al*, 1989). The HRP-coupled secondary antibodies used were: Mouse IgG HRP Linked Whole Ab - Cytiva NA931-1ML and Rabbit IgG HRP Linked Whole Ab - Cytiva NA934-1ML.

### Immuno-precipitation & slot blot

5-6×10^6^ HEK293T cells were incubated for 4 hours with VLPs carrying *d*RnhA. Cells were washed twice with cold phosphate-buffered saline (PBS), scraped in cold PBS and incubated in hypotonic buffer (20 mM Tris-HCl pH 7.4; 15 mM NaCl; 0.1% Triton X-100; 0.5% IGEPAL) for 15 minutes on ice, in the presence of protease inhibitors (cOmplete EDTA-free protease inhibitor cocktail, Roche, cat. 05056489001). The chromatin fraction was pelleted (5min, 800g), washed in hypotonic buffer, pelleted again (5min, 2000g) and solubilised for 10 min on ice in Solubilisation buffer (10 mM Tris-HCl pH 7.4; 200 mM NaCl; 0.2% Sodium deoxycholate; 0.1% SDS; 0.5% Triton X-100) in the presence of protease inhibitors. Chromatin was then sonicated to ∼600 bp fragments using a Pixul apparatus (Active Motif) with the following parameters: Pulse (50N), PRF (1 kHz), Burst rate (20 Hz), time (15min). After centrifugation (5min, 6000g), 10% of the supernatant were kept for Western blotting (INPUT sample). The rest was diluted in Dilution buffer (10 mM Tris-HCl pH 7.4; 200 mM NaCl; 0.5%Triton X-100) supplemented with protease inhibitors. *d*RnhA was captured on 50 µL V5-Trap® Magnetic Particles (Proteintech, cat. v5td) prepared according to the manufacturer’s instructions. After a one-hour incubation at 4°C on a rotating wheel, magnetic beads were collected on a magnetic rack and washed four times in Wash buffer (10 mM Tris-HCl pH 7.4; 150 mM NaCl; 0.05% IGEPAL; 0.5 mM EDTA). 10% of the beads were kept for Western blotting (IP sample). Nucleic acids were eluted from the remaining beads by a 30-minute incubation in Elution buffer (SDS 1%; NaHCO_3_ 0.1 M). Supernatants were collected into fresh tubes and digested with 200 µg of Proteinase K for 1h at 37°C. Nucleic acid were recovered using Phenol:Chloroform:Isoamyl Alcohol [25:24:1] purification and Ethanol precipitation. After extensive washes with fresh Ethanol 80°, pellets were resuspended in Tris-lowEDTA (10 mM Tris-HCl pH 8.0; 0.5 mM EDTA) and digested with 0.04 U/µL RNase III (NEB, cat. M0245L) in CutSmart Buffer (NEB, cat. B6004S) for 2h at 37°C. For negative controls, the samples were also digested with 0.05 U/µL of RNase H (NEB, cat. M0297L) in the same buffer. The different digests were deposited on a Hybond^TM^-N+ membrane (Amersham, cat. RPN203B) using a slot blot apparatus (Bio-Dot^®^ SF Microfiltration apparatus, BioRad) according to the manufacturer’s instructions. Two successive pulses with UV (0.12J) were used to crosslink nucleic acids to the membrane. After blocking the membrane with 5% milk dissolved 1X PBS containing 0.02% Tween-20, RNA:DNA hybrids and dsDNA were quantified respectively using the S9.6 antibody (lab stock) and the ab27156 (Abcam) antibody. Signals were revealed using SuperSignalTM West Femto Maximum Sensitivity Substrate (ThermoScientific, cat. 34096).

### R-loop transfection and visualization

*In vitro* transcription was used to produce R-loops at *mAirn* as previously published (Carrasco-Salas *et al*, 2019). After *in vitro* transcription, the plasmids were purified with AMPure beads (Beckman Coulter, cat. A63881; 1.8x ratio) according to the manufacturer’s instructions and then quantified using Qubit DNA high sensitivity kit (Thermo Scientific, cat. Q32851) on a Qubit 2 fluorometer (Thermo Scientific, cat. Q32866). ∼100 ng of the R-loop-containing plasmid and 2 µg of an empty carrier plasmid (pBLADE) were transfected into ∼4.10^5^ HeLa cells grown on coverslips with Lipofectamine 3000 (Invitrogen) using the manufacturer’s instructions. Three hours later, cells were incubated in fresh medium for another three hours, washed twice in 1X PBS and fixed for 15 minutes in 4% formaldehyde. After a wash in 1X PBS, cells were incubated 10 minutes in 1X PBS containing 0.2% Triton-X100 and blocked 30 minutes in 1X PBS containing 3% BSA. The S9.6 was added at 550 ng/mL and incubated overnight at 4°C. After washing cells twice in 1X PBS, cells were incubated in the dark for 30 minutes at RT with the relevant secondary antibody diluted 1/200 in 1X PBS containing 3% BSA. After washing cells twice in 1X PBS, cells were incubated in the dark for 1 hour at RT with 0.5 µg of the anti-V5 antibody already coupled to the relevant secondary antibody using the FlexAble 2.0 Coralite® Plus 488 antibody labelling kit for mouse IgG2a according the to manufacturer’s instructions. After extensive washes in 1X PBS, cells were incubated with 1 µg/mL DAPI in water for 15 minutes, washed with water and mounted on a slide. Co-localization was assessed manually by two independent authors in a single-blind manner.

### ImmunoStaining

∼2.5×10^5^ HeLa cells were plated on coverslips (Marienfield, cat. 0101050). 24h later, cells were treated as indicated, washed twice with cold PBS and pre-extracted with cold PBS-Triton 0.2% (Sigma-Aldrich, cat. T8787) for 2 min on ice. After 2 washes with PBS, cells were fixed in the dark for 15 min with freshly-made PBS containing 3% formaldehyde (thermoScientific, cat. 28908). After 2 washes with PBS, cells were permeabilized with PBS-Triton 0.2% for 5 min and then washed again twice with PBS. Cells were then blocked in a humidity chamber with PBS containing 3% BSA (PBS-BSA) for one hour, and incubated with the relevant primary antibodies in PBS-BSA overnight at 4°C. After 2 washes with PBS-BSA and 1 wash with PBS, coverslips were incubated for 30min in a humidity chamber with the relevant secondary antibodies diluted in PBS-BSA. After 2 washes with PBS, cells were incubated with 0.5 µg/mL DAPI (5min) and coverslips were mounted with VECTASHIELD® Antifade Mounting Medium (Vector Laboratories, cat. H-1000). Image acquisition was carried out using a confocal microscope LSM800 (Zeiss). The primary antibody SV5-Pk1 (BIO-RAD, cat. MCA1360) was used against V5-tagged proteins. The following secondary antibodies were used: F(ab’)2-Goat anti-Mouse IgG (H+L) Cross-Adsorbed Secondary Antibody, Alexa Fluor™ 568 (Invitrogen, cat. A11019) and F(ab’)2-Goat anti-Rabbit IgG (H+L) Cross-Adsorbed Secondary Antibody, Alexa Fluor™ 488 (Invitrogen, cat. A1070).

### Cell synchronization

HeLa cells were incubated for 18 hours with 2 mM thymidine (Sigma-Aldrich, cat. T1895). After three washes with the growth medium, cells were released into fresh medium for 200min.

### Flow cytometry

Cells were incubated with 25 µM EdU (Invitrogen, cat. C10632) for 30min. After one wash with PBS, cells were collected and fixed for 15 min in 1X PBS containing 4% formaldehyde (RT in the dark). Cells were then washed twice with PBS containing 3% BSA (PBS-BSA). To monitor nascent DNA, EdU incorporation was quantified using the Click-iT Plus EdU Flow Cytometry Assay Kit (Invitrogen, cat. C10632) following the manufacturer’s instructions. Total DNA was quantified using 1 µg/mL DAPI. Flow cytometry was carried out using a MACSQuant VYB cytometer (Miltenyi Biotec).

### Proximity Ligation Assay (PLA)

∼1.5×10^5^ HeLa cells were used per experiment. Cells were washed twice with cold PBS and then incubated for 10 min with cold PBS containing 0.5% Triton (Sigma-Aldrich, cat. T8787) and protease inhibitors (cOmplete EDTA-free protease inhibitor cocktail, Roche, cat. 05056489001). After 2 washes with PBS containing protease inhibitors, coverslips were fixed for 15 min in the dark with fresh PBS containing 4% formaldehyde (thermoScientific, cat. 28908) and protease inhibitors. After two 5-min washes with PBS, cells were permeabilized with PBS containing 0.2% Triton for 10 min and then washed twice with PBS (5 min each). Proximity Ligation Assay was carried out using NaveniFlex Cell MR Red (Navinci NC.MR.100.Red) according to the manufacturer’s instructions. Coverslips were mounted using Duolink® In Situ Mounting Medium with DAPI (Sigma-Aldrich, cat. DUO82040). Image acquisition was carried out using a confocal microscope LSM800 (Zeiss). In Figure 4, PLA signals were analysed by clustering as explained below. In Figure 5C, PLA foci were quantified automatically with the Arivis software (Zeiss).

### Clustering of PLA signals

All .czi stack files were analysed using the procedure explained here: https://gitbio.ens-lyon.fr/LBMC/RMI2/mundi_centro. Briefly, for every stack, each z slice was saved as an independent .tiff file. Nuclei in each z slice were detected on the DAPI channel using the pre-trained nuclei model of Cellpose (Stringer *et al*, 2021; Pachitariu & Stringer, 2022). ROIs for each nucleus along the z axis were reassociated using the *AgglomerativeClustering* function of scikit-learn (Pedregosa *et al*, 2011). For each nucleus, downstream analysis was carried out on the slice with the bigger area. Pixel values and ROIs properties were saved to dedicated files and a dedicated statistical analysis was used to sort nuclei into different clusters depending on the intensity of their fluorescent signals. Briefly, nuclei in all conditions and all biological replicates were pooled and analysed together in an unbiased manner. The distribution of pixel intensities in each nucleus was determined and used to classify nuclei into four different clusters using the kmeans function of scikit-learn (Pedregosa *et al*, 2011). The proportion of nuclei in each cluster was determined for each condition across biological replicates.

### DNA fibre spreading

DNA fibre spreading was performed as described previously (Jackson & Pombo, 1998; Coquel *et al*, 2018). Briefly, sub-confluent cells were sequentially labelled with 10 µM 5-iodo-2’-deoxyuridine (IdU) and with 40 µM 5-chloro-2’-deoxyuridine (CldU) for the indicated times. About 1000 cells were loaded onto a glass slide (StarFrost) and lysed with spreading buffer (200 mM Tris-HCl pH 7.5, 50 mM EDTA, 0.5% SDS) by gently stirring with a pipette tip. The slides were tilted slightly and the surface tension of the drops was disrupted with a pipette tip. The drops were allowed to run down the slides slowly, then air dried, fixed in methanol/acetic acid 3:1 for 10 minutes, and allowed to dry. Glass slides were processed for immunostaining with mouse anti-BrdU to detect IdU (clone B44, BD Biosciences, Ref: 347580), rat anti-BrdU to detect CldU (clone BU1/75, Abcam, ab6326), mouse anti-ssDNA antibodies and corresponding secondary antibodies conjugated to various Alexa Fluor dyes. Nascent DNA fibres were visualized using immunofluorescence microscopy (Zeiss ApoTome). The acquired DNA fibre images were analysed by using MetaMorph Microscopy Automation and Image Analysis Software (Molecular Devices) and statistical analysis was performed with GraphPad Prism (GraphPad Software). The lengths of at least 150 IdU/CldU tracks were measured per sample.

### Slot blot to quantify RNA:DNA hybrids

After two washes with 1X PBS, 1-2×10^6^ cells grown in 6 cm plates were lysed directly on plates with 1 mL of lysis buffer (10 mM Tris pH 8.0, 1% SDS, 600 µg/mL Proteinase K, 20 mM EDTA pH 8.0). The lysate was then scraped off the plate and transferred into a 1.5 mL tube and incubated at 55°C for 4 hours. Purification with Phenol:Chloroform:Isoamyl Alcohol [25:24:1] pH8.0 was carried out in MaXtract High Density tubes (Qiagen, cat. 129065) and nucleic acids were precipitated with 2.5 volumes of Ethanol. Extensive washes with fresh Ethanol 80° were carried out before nucleic acids were resuspended in 10 mM Tris pH 8.0. The concentrations of genomic DNA (gDNA) were determined by quantitative PCR (qPCR) using primers against Alu elements (Alu F: GTGGCTCACGCCTGTAATC & Alu R: CAGGCTGGAGTGCAGTGG). 4 µg of gDNA was then digested with HindIII-HF (NEB, cat. R3104S) and EcoRI-HF (NEB, cat. R3101S) according to the manufacturer’s instructions. For negative controls, 5U/µg of RNase H (NEB, cat. M0297L) was added to the digest. Different amounts of gDNA were deposited on a Hybond^TM^-N+ membrane (Amersham) using a slot blot apparatus (Bio-Dot^®^ SF Microfiltration apparatus, BioRad) according to the manufacturer’s instructions. Two successive pulses with UV (0.12J) were used to crosslink the DNA to the membrane. After blocking the membrane with 5% milk dissolved 1X PBS containing 0.02% Tween-20, RNA:DNA hybrids and dsDNA were quantified respectively using the S9.6 antibody (lab stock) and the ab27156 (Abcam) antibody. Alternatively, dsDNA was revealed with Methylene blue. The concentrations of genomic DNA (gDNA) in the last dilution of gDNA were determined by quantitative PCR (qPCR) using primers against Alu elements (see above) to confirm that an equal amount of gDNA was deposited on a membrane in all samples.

### DRIP

gDNA was extracted and quantified as above. It was then sonicated using a Pixul apparatus (Active Motif) with the following parameters: Pulse (50N), PRF (1 kHz), Burst rate (20 Hz), time (35min). For negative controls, gDNA was digested for at least 5 hours at 37°C with 5U/µg RNase H in the buffer provided by the manufacturer (NEB, cat. M0297L). 4 or 20 µg of sonicated gDNA was used for DRIP-qPCR or DRIP-seq respectively. The S9.6 antibody (lab stock) was coupled to Dynabeads^TM^-Protein A (Invitrogen, cat. 10002D) according to the manufacturer’s instructions. 2% of gDNA was set aside (input). The rest of the gDNA and the S9.6 antibody were mixed in cold IP buffer (10 mM Tris pH8.0, NaCl 140 mM, 0.5% Triton, 1X cOmplete EDTA-free protease inhibitor cocktail, (Roche, cat. 05056489001) using a 2:1 ratio (by mass) and rotated for 90 minutes at 4°C. The immunoprecipitated fraction was then washed twice in IP buffer, once with Wash III buffer (10 mM Tris pH 8.0, 1 mM EDTA pH 8.0, 1% IGEPAL, 1% Sodium deoxycholate, 250 mM LiCl) and twice with TE buffer (10 mM Tris, 1 mM EDTA pH 8.0). The immunoprecipitated gDNA fragments were eluted from the beads in Elution buffer (Tris 50 mM, EDTA 10 mM, SDS 1%, 400 µg/mL Proteinase K) at 55°C for 45’ and then purified using the ChIP DNA Clean & Concentrator kit (Zymo Research) according to the manufacturer’s instructions.

### Strand-specific sequencing libraries

DRIP DNA was treated with 10 µg of RNase A (30’, 37°C) then purified on AMPure beads (Beckman Coulter, cat. A63881; 1.8x ratio) according to the manufacturer’s instructions. ∼1 ng of DNA was used to prepare sequencing libraries with the xGen™ ssDNA & Low-Input DNA Library Preparation Kit (IDT), according to the manufacturer’s instructions. Sequencing libraries were cleaned twice with AMPure beads (0.85x ratio) according to the manufacturer’s instructions and then quantified using the Qubit DNA high sensitivity kit (Thermo Scientific, cat. Q32851) on a Qubit 2 fluorometer (Thermo Scientific, cat. Q32866). Libraries were paired-end sequenced (2×150 bp) on an Illumina Novaseq6000 platform by Novogene UK.

### DRIP-seq analysis

Paired-end reads were processed, aligned on the GRCh38 genome and calibrated using the strategy described at https://gitbio.ens-lyon.fr/LBMC/Bernard/chipseq/-/tree/dev_accel_1splus?ref_type=heads, using the ‘--keep_multi_map’ option. Reads mapping to the nuclear genome were calibrated over the reads mapping to the mitochondrial genome. Peak calling was carried out with MACS2 with the following options: ‘--bdg --SPMR --nomodel -f BAMPE--min-length 300 --keep-dup all --broad’. Subsequent analyses were carried out with Deeptools on the annotated peaks whose q-value was ≥ 3 and whose length was ≥ 600 bp.

### Cut&Tag

Cut&Tag was carried out with the CUT&Tag-IT® R-loop Assay Kit according to the manufacturer’s instructions (Active Motif, cat. 53167). Libraries were quantified using the Qubit DNA high sensitivity kit (Thermo Scientific, cat. Q32851) on a Qubit 2 fluorometer (Thermo Scientific, cat. Q32866) and paired-end sequenced (2×150 bp) on an Illumina Novaseq6000 platform by Novogene UK. Paired-end reads were processed and aligned on the GRCh38 genome using the nf-core cutandrun 3.1 pipeline (https://nf-co.re/cutandrun/) (Ewels *et al*, 2020) using the cpm normalisation mode.

### RNAseq

Total RNAs were extracted with Trizol (Invitrogen™, cat. 15596026) using the manufacturer’s instructions. To extract chromatin-associated RNAs, cells were first incubated for 15’ at 4°C in Nuclei buffer (Tris 20 mM, NaCl 15 mM, Triton 0.1%, NP-40 0.5%). Nuclei were collected by a 5’ centrifugation at 800 g (4°C). The pellet was washed twice in nuclei buffer and chromatin-associated RNAs were extracted with Trizol (Invitrogen™, cat. 15596026) using the manufacturer’s instructions. For both total RNAs and chromatin-associated RNAs, residual genomic DNA was digested with the TURBO DNA-free™ kit (Invitrogen, cat. AM1907), following the manufacturer’s protocol. RNA quality and concentrations were determined using the Agilent High Sensitivity RNA ScreenTape system (cat. 5067-5579), following the manufacturer’s protocol. Sequencing libraries and paired-end sequencing (2×150bp) was performed by Novogene UK. Paired-end reads were processed and aligned on the GRCh38 genome using the nf-core rna-seq 3.9 pipeline (https://nf-co.re/rnaseq/) (Ewels *et al*, 2020). After filtering out genes whose CPM < 0.1, differential analysis with DESeq2 and scatterplot production (padj < 0.01; foldChange 1.5) was carried out using DEBrowser v1.26.3 (http://nasqar2.abudhabi.nyu.edu/DEBrowser/).

### ChIP

cells were collected with Trypsin, washed with 1X PBS and resuspended in 10 mL of fresh 1X PBS containing 1% of fresh formaldehyde. After a 10-minute incubation in the dark, glycine was added (125 mM final) and incubated for 5 minutes. Cells were washed in 1X PBS containing protease inhibitors (cOmplete EDTA-free protease inhibitor cocktail, Roche, cat. 05056489001) and the cell pellet was snap frozen at -80°C. Depending on the experiment, two different sonication procedures were implemented (see below).

- *Sonication with a Covaris apparatus (Fig 3A, B and Fig 7A):* cells pellets were resuspended in 1 mL of Lysis buffer (10 mM Tris-HCl pH 7.5; 140 mM NaCl; 0.5% IGEPAL) containing protease inhibitors. Cells were then sonicated using a Covaris S220 apparatus (2 min; power: 75; duty factor: 2; cycle/burst: 200) and pelleted (5’, 900 g, 4°C). Pellets were rinsed once in 1 mL Lysis buffer containing protease inhibitors and resuspended in 1 mL of Shearing buffer (10 mM Tris-HCl pH7.5; 140 mM NaCl; 1 mM EDTA pH8.0; 0.5% IGEPAL; 0.1% SDS) containing protease inhibitors before a second round of sonication with a Covaris S220 apparatus (15 min; power: 140; duty factor: 5; cycle/burst: 200). After clarification (10 min, 19000 g, 4°C; twice), 50 µL of the lysate was kept as Input, 100 µL was used to analyse the shearing efficiency and 700 µL was incubated (overnight, 4°C) with 50 µL of Protein A-Dynabeads coupled to 10 µg of V5 antibody according to the manufacturer’s instructions.
- *Sonication with a Pixul apparatus (Fig 3C):* cells pellets were resuspended at the concentration of 10^6^ cells/100 µL in Shearing buffer (10 mM Tris-HCl pH 7.5; 140 mM NaCl; 0.5% IGEPAL; 1 mM EDTA pH 8.0; 0.1% SDS) containing protease inhibitors and sonicated in plates using a Pixul Sonicator (50 min; burst rate: 20Hz; Pulse: 50N; PRF: 1kHz). The volume of sample was completed with shearing buffer to reach 1 mL, and the sample was then clarified by centrifugation (10 min, 19000 g, 4°C; twice). After clarification, 50 µL of the lysate was kept as Input, 100 µL was used to analyse the shearing efficiency and 700 µL was incubated (overnight, 4°C) with 50 µL of Protein A-Dynabeads coupled to 10 µg of V5 antibody according to the manufacturer’s instructions.

The rest of the protocol did not differ between the two sonication procedures. The beads were washed successively with 1 mL of wash buffer #1 (20 mM Tris pH7.5; 150 mM NaCl; 2 mM EDTA pH8.0; 1% Triton; 0.1% SDS), 1 mL wash buffer #2 (20 mM Tris pH7.5; 500 mM NaCl; 2 mM EDTA pH8.0; 1% Triton; 0.1% SDS), 1 mL wash buffer #3 (10 mM Tris pH7.5; 2 mM EDTA pH8.0; 1% Na-deoxycholate; 1% IGEPAL; 250 mM LiCl; 5’ RT with rotation) and then twice with 1 mL TE pH8.0. Beads were resuspended in 100 µL of Elution buffer (10 mM Tris pH7.5; 1 mM EDTA pH8.0; 1% SDS) containing 140 µg of Proteinase K and incubated for 4 hours at 65°C. The supernatant was then purified using the ChIP DNA Clean & Concentrator kit (Zymo Research) according to the manufacturer’s instructions.

## ACKNOWLEDGMENTS

We are extremely grateful to Benoit Palancade and Aurèle Piazza for their comments on the manuscript. We thank Jana Dobrovolná and Pavel Janscak for providing the U2OS cell line over-expressing hRNH1 fused to GFP and to Els Verhoeyen for the gift of pBaEVRless. This study was supported by funding from Fondation pour la Recherche Médicale – FRM attributed to AB (contrat doctoral #ECO202306017384), from Ligue contre le Cancer (comité du Rhône) attributed to VV and EPR (Appel d’offres régional 2021), from Fondation ARC pour la recherche sur le cancer attributed to VV (ARCPJA2021060003952), from Agence Nationale de la Recherche (ANR) attributed to PP and VV (project ANR-19-CE12-0016-04 and project ANR-23-CE12-0020-03), from Labex Ecofect (ANR-11-LABX-0048) of the Université de Lyon, within the program Investissements d’Avenir (ANR-11-IDEX-0007) and from the European Research Council (ERC-StG-LS6-805500) attributed to EPR under the European Union’s Horizon 2020 research and innovation programs. We acknowledge the contribution of the LBMC Bioinformatics Hub and in particular the help of Laurent Modolo with the building of the DRIP analysis pipeline. We gratefully acknowledge support from the CBPsmn (PSMN, Pôle Scientifique de Modélisation Numérique) of the ENS de Lyon for computing resources. We acknowledge the contribution of SFR Biosciences (Université Claude Bernard Lyon 1, CNRS UAR3444, Inserm US8, ENS de Lyon) and the help of the staff of LyMIC-PLATIM, especially Elodie Chatre, for assistance with confocal microscopy and of Véronique Barateau for assistance with flow cytometry. We thank the staff of CIQLE, especially Bruno Chapuis, for their assistance with the Arivis software.

## DISCLOSURE AND COMPETING INTERESTS STATEMENT

The authors declare that they have no conflict of interest.

## DATA AVAILABILITY

### Lead contact

Further information and requests for resources and reagents should be directed to and will be fulfilled by the lead contact, Vincent Vanoosthuyse.

### Materials availability

Plasmids generated in this study are available upon request.

### Data and code availability

- The raw Illumina paired-end sequencing data and the processed files in a bigwig format have been deposited to NCBI Gene Expression Omnibus (GEO) under accession GSE288898, GSE288899, GSE288900 and are publicly available from the date of publication.
- All original code has been deposited to Github as described in the methods section and are publicly available from the date of publication.

## EXPANDED VIEW FIGURE LEGENDS

**Figure EV1:**
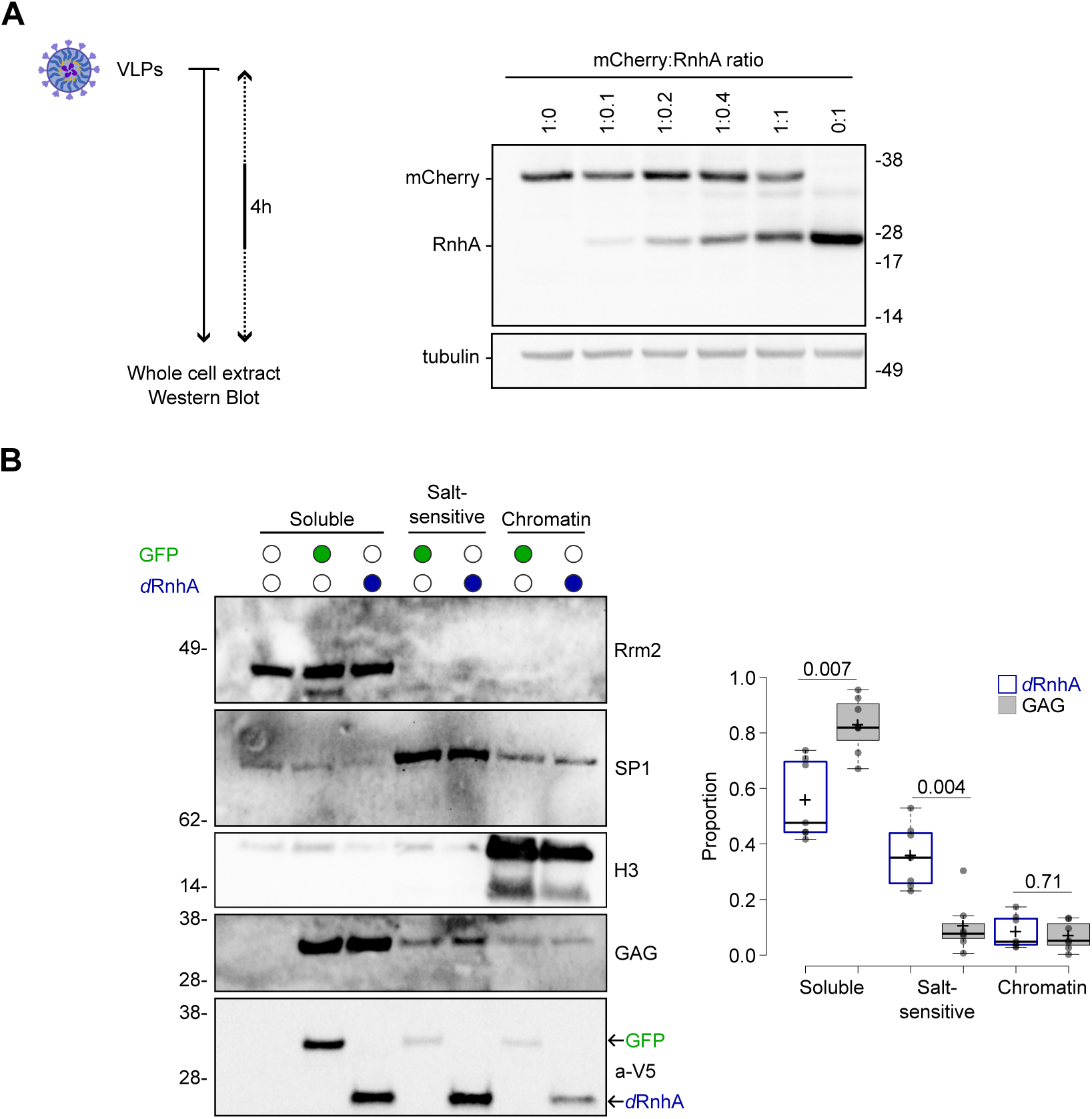
*iCoRD*-delivered RnhA associates with chromatin. All experiments were carried out in HeLa cells. **A.** Western blot analysis of total protein extracts of cells treated with VLPs co-delivering different amounts of V5-tagged mCherry and RnhA. Tubulin was used as loading control. **B.** Cells were incubated with VLPs delivering either GFP or RnhA^D10N^ (*d*RnhA) for four hours. Protein extracts were fractionated into three fractions: (i) soluble, (ii) salt-sensitive association with chromatin and (iii) tight association with chromatin. Rrm2, SP1 and Histone H3 were used as markers for the different fractions. The retroviral protein GAG and GFP remained mainly in the soluble fraction, whilst a fraction of *d*RnhA associated with chromatin in a salt-sensitive manner. (left) a typical experiment; (right) quantifications of 7 experiments (p-value determined from a Wilcoxon-Mann-Whitney test).

**Figure EV2:**
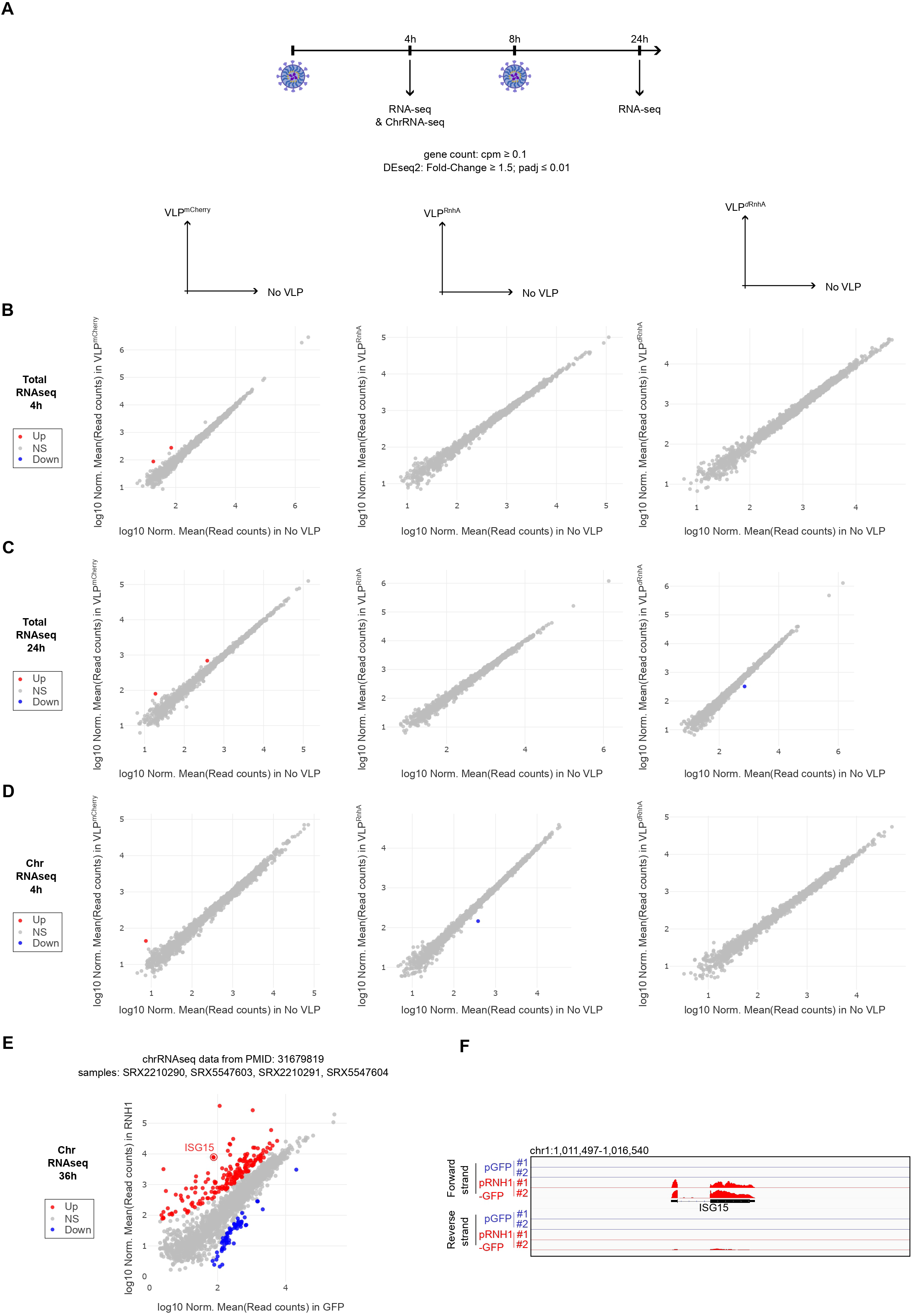
*iCoRD*-delivered RnhA does not impact the steady state transcriptome. All experiments were carried out in HeLa cells. **A.** Schematic representation of the experiment. **BCD.** Scatterplot comparisons of the transcriptomes of cells treated or not with VLPs delivering mCherry (left), RnhA (center) or *d*RnhA (right). Total RNAs were analysed after a 4-hour (**B.**) or a 24-hour (**C.**) incubation. In **D.**, chromatin-associated RNAs were analysed after a 4-hour incubation. **EF.** Re-analysis of the indicated published RNA-seq data (GSM2335051, GSM3681017, GSM2335052, GSM3681018) (Tan-Wong *et al*, 2019) comparing the chromatin-associated RNAs of cells over-expressing GFP or GFP fused to human RNase H1 for 36 hours. **E.** Scatterplot comparisons. ISG15 is indicated on the plot. **F.** RNA-seq signals around ISG15 (published data from (Tan-Wong *et al*, 2019)).

**Figure EV3:**
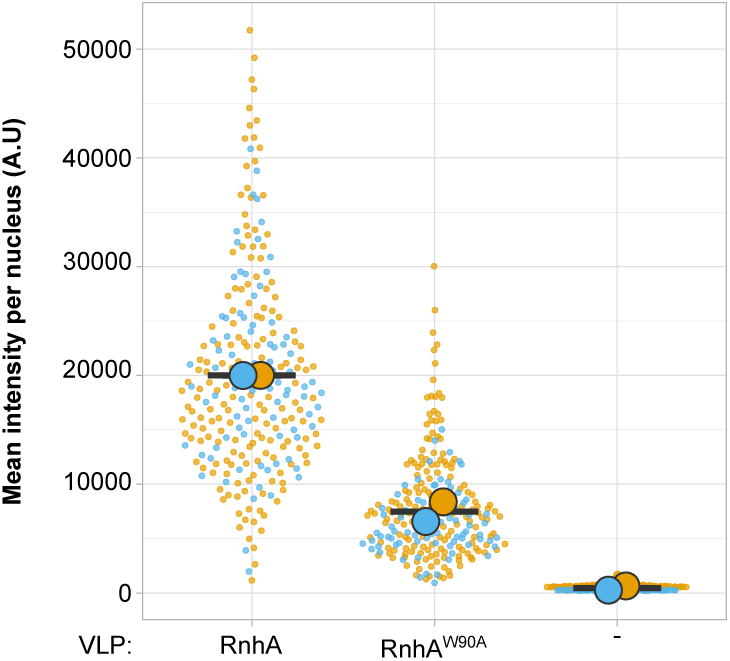
The RnhA^W90A^ mutant is less efficiently delivered by VLPs than RnhA. HeLa cells were incubated for four hours with VLPs delivering either RnhA or RnhA^W90A^. Immuno-staining was used to quantify the nuclear levels of VLP-delivered RnhA or RnhA^W90A^ in cells. The distribution of intensities in individual nuclei is shown. The circles of different colours indicate the average nuclear intensity in 2 independent experiments.

**Figure EV4:**
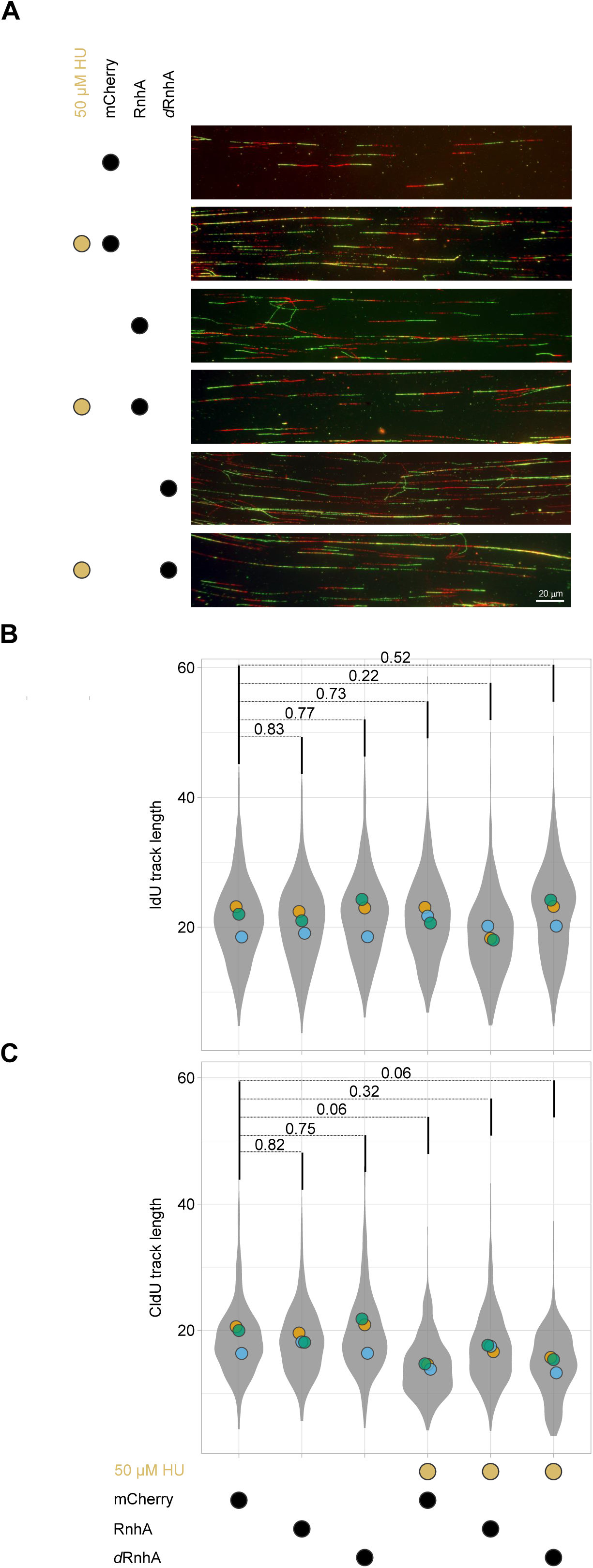
Quality controls for the chromatin spreading experiments (related to Figure 4A). After short pulses with IdU and CldU, chromatin spreading was used to monitor replication fork progression. **A.** Examples of chromatin fibres. **BC.** The distribution of values for individual track length for IdU **(B)** and CldU **(C)** is shown as superplots. The distribution of values is shown in grey and the circles of different colours indicate the mean values in 3 independent experiments. Comparisons between the means of the replicates were carried out and Welch’s t-test was performed to calculate the indicated p-values.

**Figure EV5:**
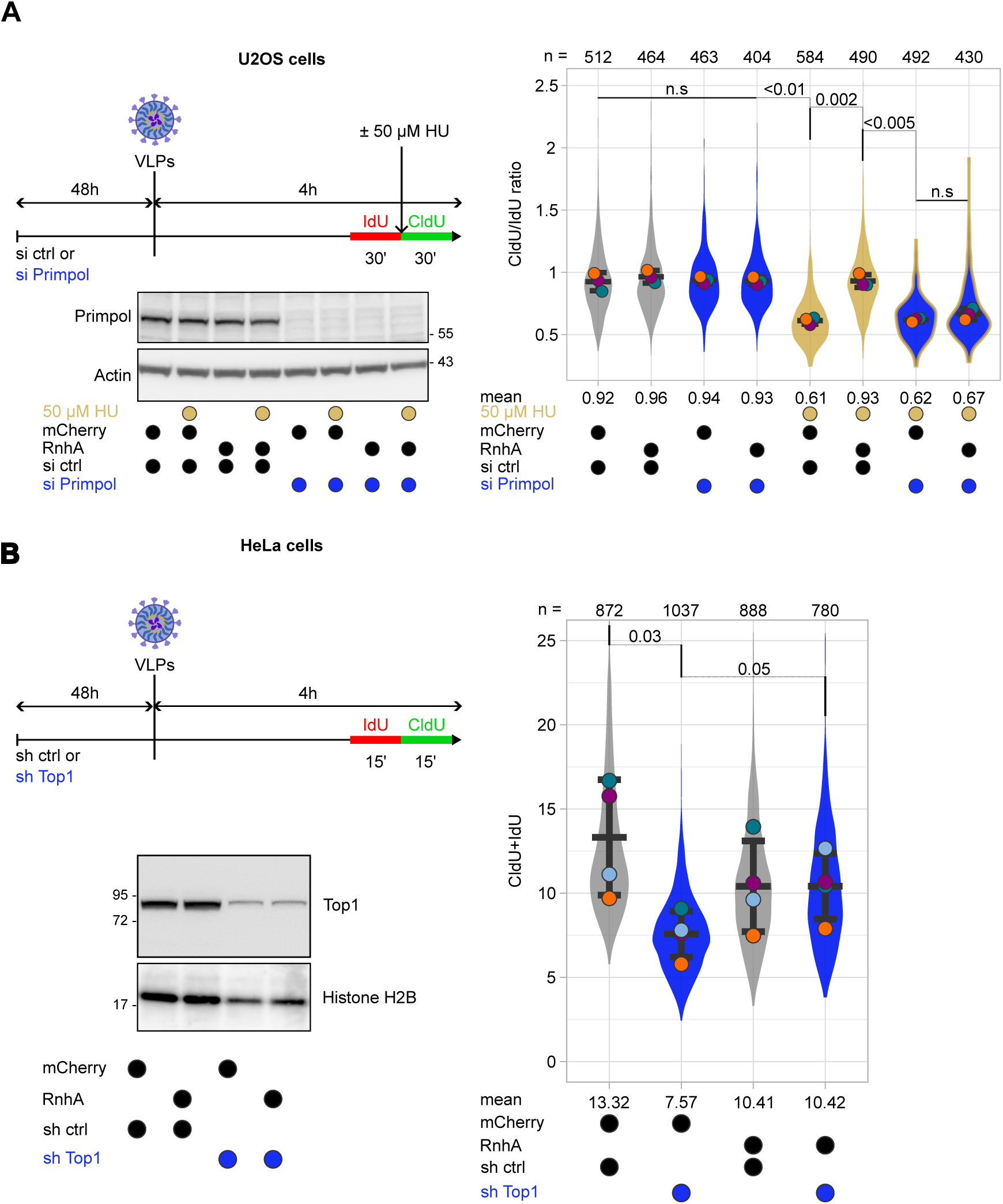
*iCoRD*-delivered RnhA rescues the progression of stressed replication forks (related to Figure 5). The cell line used and the timeline of the experiment is shown for each panel. After short pulses of the indicated duration with IdU and CldU, chromatin spreading was used to monitor replication fork progression. Measurements for individual fibres are represented on superplots, representing the distribution of values. The circles of different colours indicate the mean values for individual experiments. The total number of chromatin fibres that were scored is indicated at the top of graphs. Comparisons between the means of the replicates were carried out and Welch’s t-test was performed to calculate the indicated p-values. **(A**) Primpol is necessary for the rescue of replication fork progression by *iCoRD*-delivered RnhA in cells treated with 50 µM HU. U2OS cells were depleted or not with Primpol and treated or not with 50 µM HU for 30 mins. Western blot was used to confirm the depletion of Primpol. Actin was used as loading control. Superplots represent the CldU/IdU ratio in 3 independent experiments. (**B**). *iCoRD*-delivered RnhA rescues the progression of replication forks upon depletion of Top1. HeLa cells were depleted or not for Top1. Western blot was used to confirm the depletion of Top1. Histone H2B was used as loading control. Superplots represent the sum of CldU and IdU track lengths in 4 individual experiments.

**Figure EV6:**
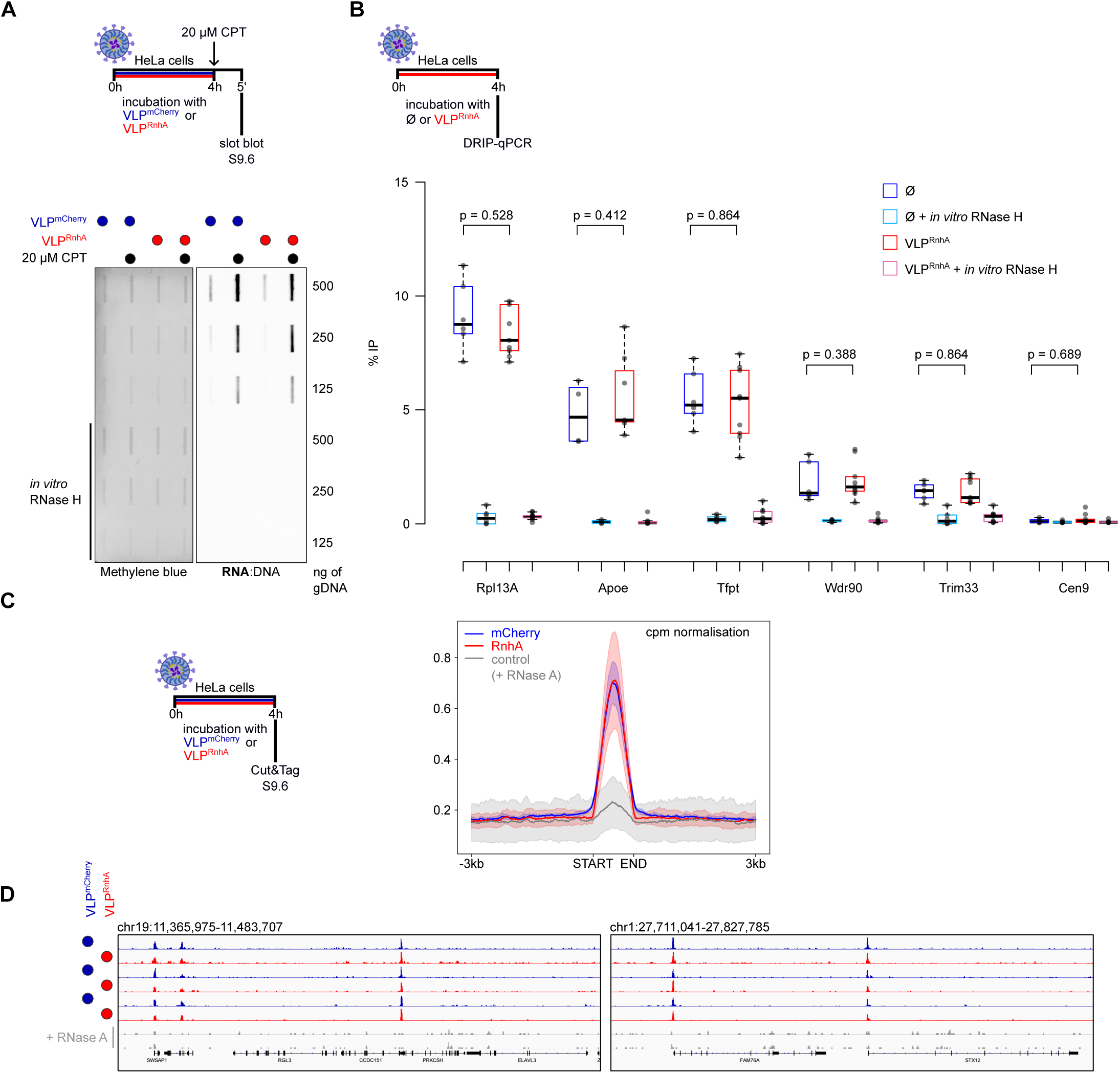
VLP-delivered RnhA does not impact DRIP or Cut&Tag signals. **A.** The experiment was carried out in HeLa cells. (top) Timeline of the experiment. (bottom) Representative slot blot analysis of total RNA:DNA hybrid levels upon a 4-hour treatment with mCherry- or RnhA-containing VLPs, after cells were treated or not with 20 µM CPT for 5 mins. Methylene blue was used to label DNA. The *<u>in vitro</u>* RNase H treatment confirms that the S9.6 signals correspond to RNA:DNA hybrids. **B.** The experiment was carried out in HeLa cells. (top) Timeline of the experiment. (bottom) DRIP-qPCR signals at different loci cells incubated or not (Ø) with RnhA-containing VLPs for four hours. The *<u>in vitro</u>*RNase H treatment confirms that DRIP signals correspond to RNA:DNA hybrids. **CD.** Cut&Tag with the S9.6 antibody was carried out to map R-loops after HeLa cells were incubated with either GFP-containing or RnhA-containing VLPs for four hours (n = 3). **C.** Cpm-normalised signals around called peaks. **D.** Examples of Cut&Tag signals.

**Figure EV7:**
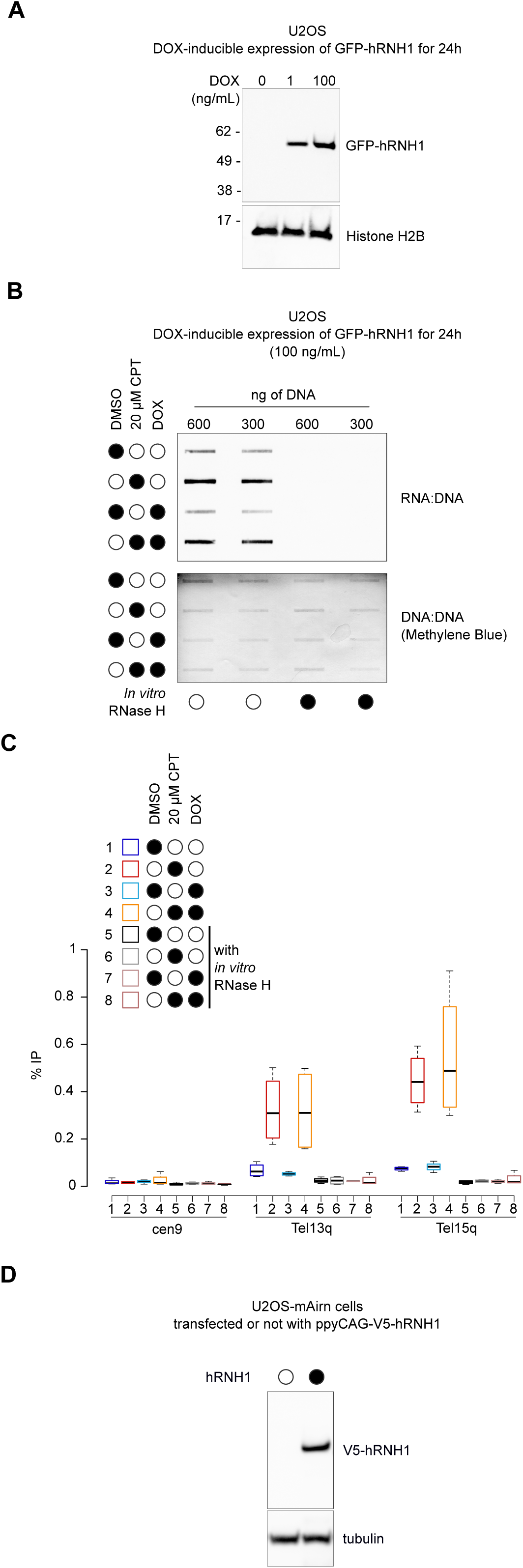
The over-expression of human RNase H1 (hRNH1) does not impact DRIP signals. AC. The experiments were carried out in U2OS cells stably expressing GFP-hRNH1 (Teloni *et al*, 2019). **A.** Western blot of chromatin-associated proteins showing the dose-dependent induction of GFP-hRNH1 by the addition of doxycycline (DOX). Histone H2B was used as loading control. **BC.** The over-expression of GFP-hRNH1 was induced or not by addition of 100 ng/mL of doxycycline for 24 hours. Cells were then treated for 5 minutes with 20 µM CPT or DMSO as control. RNA:DNA hybrid levels were quantified either by slot blotting (**B**) or by DRIP-qPCR (**C**) (n = 3). **D**. Western blot of total protein extracts showing the expression of V5-hRNH1 in U2OS cells over-expressing *mAirn*. Tubulin was used as loading control.

**Table EV1:**
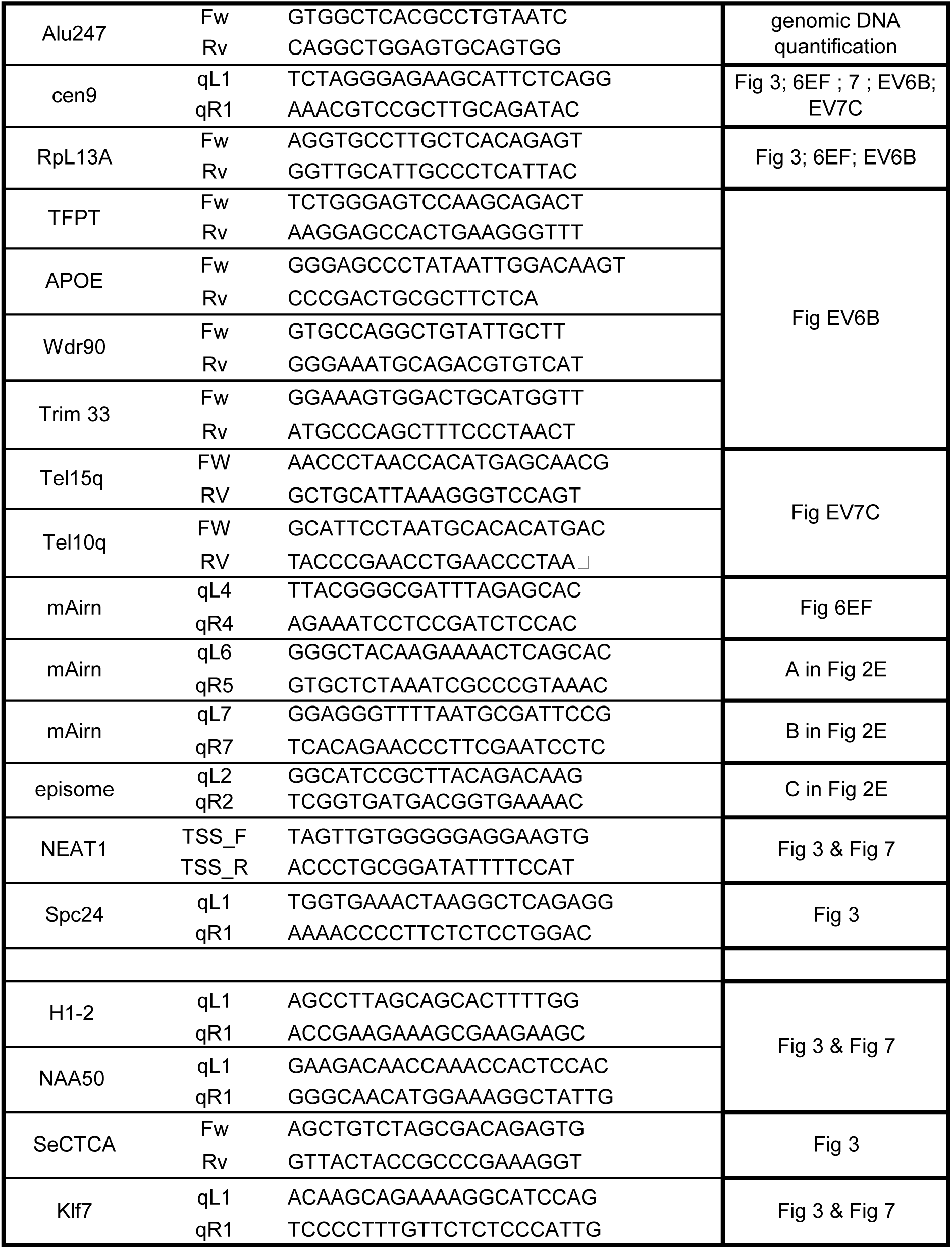
qPCR primers used in this study.

**Table EV2:**
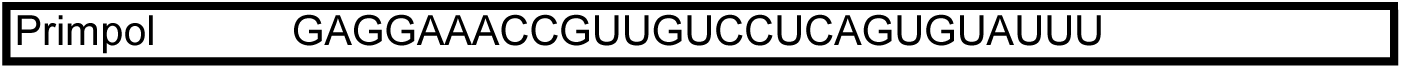
siRNA used in this study.

## Notes

### Competing Interest Statement

The authors have declared no competing interest.

